# Generating, modeling, and evaluating a large-scale set of CRISPR/Cas9 off-target sites with bulges

**DOI:** 10.1101/2023.11.01.565099

**Authors:** Ofir Yaish, Yaron Orenstein

## Abstract

The CRISPR/Cas9 system is a highly accurate geneediting technique, but it can also lead to unintended off-target sites (OTS). Consequently, many high-throughput assays have been developed to measure OTS in a genome-wide manner, and their data was used to train machine-learning models to predict OTS. However, these models are inaccurate when considering OTS with bulges due to limited data compared to OTS without bulges. Recently, CHANGE-seq, a new *in vitro* technique to detect OTS, was used to produce a dataset of unprecedented scale and quality. In addition, the same study produced *in cellula* GUIDE-seq experiments, but none of these experiments included bulges. Here, we generated the most comprehensive GUIDE-seq dataset with bulges, and trained and evaluated state-of-the-art machine-learning models that consider OTS with bulges. We first reprocessed the publicly available experimental raw data of the CHANGE-seq study to generate 20 new GUIDE-seq experiments, and hundreds of OTS with bulges among the original and new GUIDE-seq experiments. We then trained multiple machine-learning models, and demonstrated their state-of-the-art performance both *in vitro* and *in cellula* overall and when focusing on OTS with bulges. Last, we visualized the key features learned by our models on OTS with bulges in a unique representation.

**Graphical abstract:** 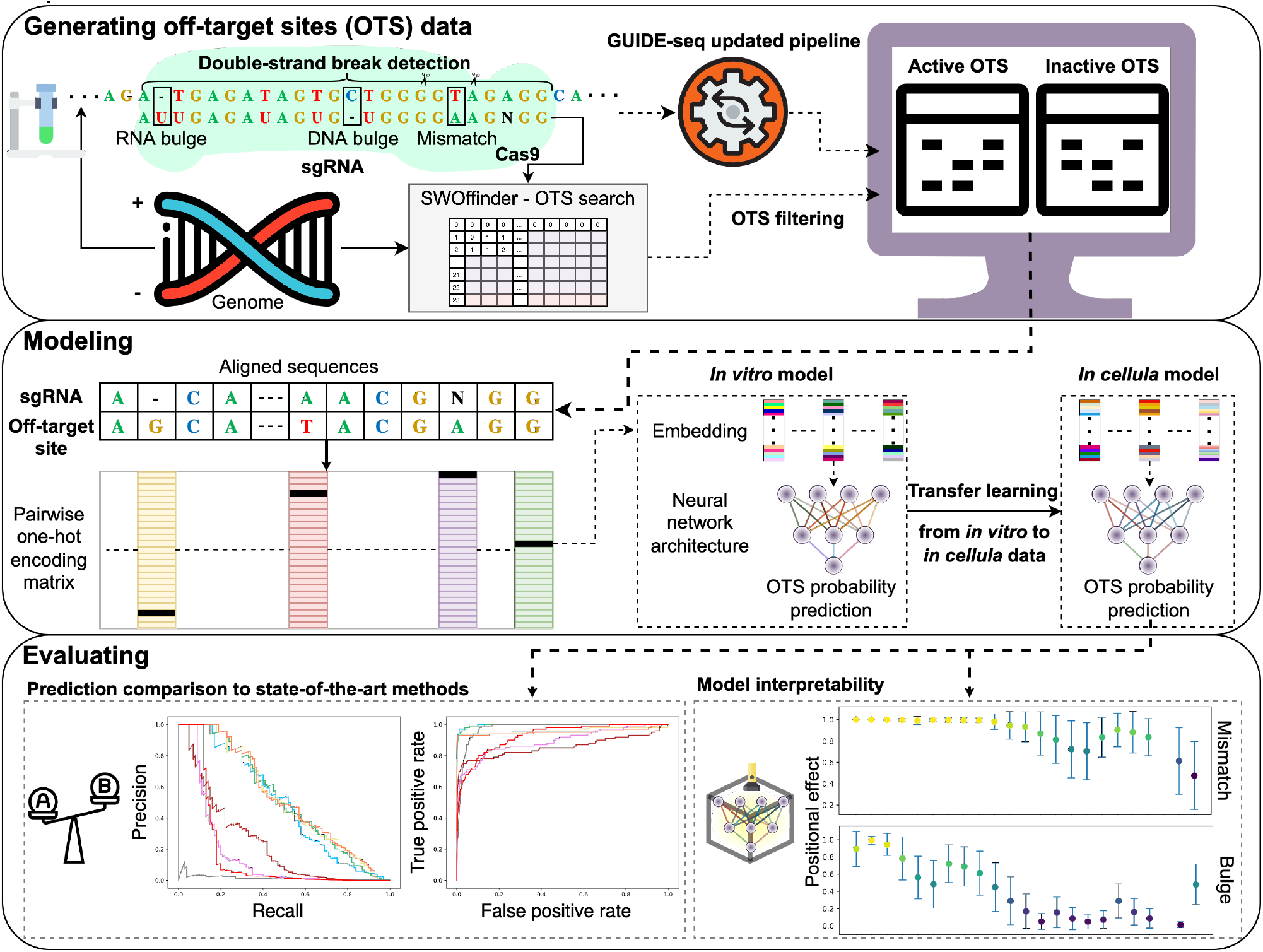

## Supplementary Note 1: Introduction

Clustered regularly interspaced short palindromic repeats (CRISPR)/Cas9 system is the preeminent geneediting technology in recent years due to its low cost, high efficiency and effectiveness, and ease of use (1; 2; 3; 4; 5). This system is composed of a Cas9 protein and an associated single guide RNA (sgRNA). The sgRNA guides the Cas9 to a target DNA sequence, and a double-strand break in the specific region is executed by the Cas9. The recognition of the DNA sequence is done via a complementarity of a 20-nucleotide (nt) sequence within the sgRNA to the genomic target upstream of a 3-nt protospacer adjacent motif (PAM) at its 3’ end. The CRISPR system is widely used in interrogating gene functions, and has wide applications in clinical detection, gene therapy, and agricultural improvement (6; 7; 8; 9; 10; 11).

Although the CRISPR/Cas9 system has many benefits, numerous studies have also shown that the mismatches between the designed sgRNA and DNA can be tolerated, resulting in the cleavage of unplanned genomic sites, termed off-target sites (OTS). Detecting OTS in CRISPR gene editing is highly important due to their potential disruptive effect, and designing sgRNAs with few OTS is desired (12; 13; 14). Many experimental techniques were developed to detect OTS, including GUIDE-seq, Digenome-Seq, HTGTS, SITE-Seq, CIRCLE-Seq, CHANGE-seq, Nuclea-seq, and BLISS (15; 16; 17; 18; 19; 20). However, these experimental techniques are limited in their ability to detect OTS with bulges, due to either experimental limits or inefficient post-processing pipelines.

Due to these and other experimental challenges, various computational methods have been developed to predict OTS based on experimental data. These methods can be split into three categories: alignment-based, ruled-based, and data-driven-based. Many data-driven methods are based on machine learning, including CRISTA, DeepCrispr, and Elevation, among many others (21; 22; 23; 24; 25; 26; 27; 28). Several methods consider bulges, including CRISPR-Net and CRISPR-IP (26; 29). However, these methods are inaccurate in predicting OTS with bulges due to limited data.

Recently, new datasets of unprecedented scale and quality were produced for both OTS *in vitro* and *in cellula* (30). The CHANGE-seq dataset contains 202, 040 unique *in vitro* OTS across 110 sgRNAs, while the GUIDE-seq dataset contains 1, 208 unique *in cellula* OTS across 58 sgRNAs, which are a subset of the 110 sgRNAs in the CHANGE-seq dataset. But, while the CHANGE-seq dataset includes 23, 692 OTS with bulges, the GUIDE-seq datasets do not include any OTS with bulges. This critical bottleneck in the number of GUIDE-seq OTS with bulges prohibits the training of any machine-learning model to predict OTS *in cellula*.

To close this gap, in this study, we generated the most comprehensive dataset of OTS with bulges to date. We reprocessed the raw experimental GUIDE-seq data that is publicly available through the CHANGE-seq study.

Through this processing, we generated more than 450 novel OTS with bulges, and 20 new GUIDE-seq experiments. Using the CHANGE-seq and GUIDE-seq datasets, and our newly generated GUIDE-seq experiments, we trained multiple machine-learning models to predict OTS with bulges. We evaluated these models on multiple datasets and demonstrated how they out-perform the state-of-the-art by utilizing transfer learning from *in vitro* to *in cellula* data. Based on the trained embedding layer, we produced novel visualizations of OTS representations with bulges, which recapitulate known and novel CRISPR/Cas9 biology.

## Materials and Methods

### A. Datasets and our updated preprocessing

#### CHANGE-seq and GUIDE-seq datasets

We used *in vitro* and *in cellula* high-throughput CRISPR OTS datasets (Table 1). CHANGE-seq is an advanced scalable, highly sensitive, and unbiased assay for measuring the genome-wide activity of CRISPR/Cas9 nucleases *in vitro* (30). The developers of CHANGE-seq applied their method to 110 sgRNA targets with an NGG PAM sequence across human primary T cells. They processed the experimental output to identify OTS with up to 1 DNA or RNA bulge and 4 mismatches compared to the sgRNA, or up to 6 mismatches and no bulges, which resulted in 202, 040 OTS.

**Table 1.** Datasets summary. Details of our processed off-target sites (OTS) datasets used for training and testing our machine-learning models

| Dataset | sgRNAs | Total active OTS | Active OTS with bulges*<br>(RNA / DNA) | Total inactive OTS | Inactive OTS with bulges*<br>(RNA / DNA) | Notes |
| --- | --- | --- | --- | --- | --- | --- |
| CHANGE-seq | 110 | 202,040 | 23,692<br>(17,267 / 6,425) | 4,734,238 | 1,725,650<br>(1,391,017 / 334,633) | Active sites as obtained in the CHANGE-seq study (30) |
| GUIDE-seq | 58 | 1,520 | 312<br>(212 / 100) | 2,118,820 | 739,258<br>(582,234 / 157,024) | 1. Active sites with bulges obtained using our updated raw data reprocessing<br>2. Duplicate active sites were aggregated |
| New GUIDE-seq experiments | 20 | 646 | 147<br>(100 / 47) | 1,150,063 | 438,228<br>(360,471 / 77,757) | New experiments obtained using our updated raw data reprocessing |
| Tsai 2015 GUIDE-seq | 10 | 420 | 80<br>(57 / 23) | 1,934,789 | 752,275<br>(655,131 / 97,144) | Active sites with bulges obtained using our updated raw data reprocessing |
| Chen 2017 GUIDE-seq | 6 | 205 | 51<br>(36 / 15) | 1,741,443 | 686,407<br>(605,123 / 81,284) |  |
| Listgarten 2018 GUIDE-seq | 23 | 86 | 11<br>(8 / 3) | 579,009 | 180,607<br>(136,953 / 43,654) |  |
| Refined TrueOT | 10 | 100 | 27<br>(20 / 7) | 354,252 | 121,012<br>(90,792 / 30,220) | Among experimentally validated sites, sites with an editing rate greater than 0.1% are considered active sites (39; 40) |

\*OTS with up to 1 DNA or RNA bulge and 4 mismatches. Without bulge, sites with up to 6 mismatches are allowed.

GUIDE-seq is an *in cellula* and unbiased approach for global detection of DNA double-stranded breaks introduced by CRISPR RNA-guided-nucleases (15). GUIDE-seq measurements are affected by chromatin accessibility and other epigenetic and cellular factors (31). The developers of CHANGE-seq conducted GUIDE-seq experiments to compare their measurements with the measurements produced by the CHANGE-seq assay.

Due to an outdated data processing pipeline, the published GUIDE-seq dataset lacks OTS with bulges. To address this gap, we applied an updated version of the GUIDE-seq pipeline (available at github.com/tsailabSJ/guideseq) to the raw data, which we downloaded from NCBI Sequence Read Archive (SRA) PR-JNA625995. We identified 1, 520 unique OTS over 58 sgRNAs across the hg38 reference genome, including 312 sites containing DNA or RNA bulges, where we set the same bulge and mismatches thresholds as were applied on the CHANGE-seq data. To remove duplicate OTS from the data, we averaged the read count of non-unique OTS over their replicates.

#### Newly generated GUIDE-seq experiments

To apply the updated version of the GUIDE-seq pipeline, we had to acquire the raw high-throughput sequencing (HTS) data files along with their corresponding DNA barcodes. While the sequencing data of the GUIDE-seq experiments is publicly available through NCBI SRA, the DNA barcodes were not published. Upon request, we obtained the sequencing barcode files from the researchers of the CHANGE-seq study who conducted and processed these GUIDE-seq experiments. However, we encountered inconsistencies between some of the barcode files and their raw sequencing data files. To address these inconsistencies, we applied the GUIDE-seq pipeline using all available barcodes on each raw sequencing data file. While this process is computationally intensive, it confirms which barcodes are associated with each raw sequencing data file based on the demultiplexing step of the pipeline. We set the minimum number of reads for the demultiplexing step to the default value of 10, 000. Due to the unexpectedly large number of demultiplexed files (more than 58 experiments and their replicates), we suspected that the raw data contained additional GUIDE-seq experiments. To investigate further, we examined if any of the 110 sgRNAs of the CHANGE-seq sgRNA library had one of the highest read counts in the sequencing data. In addition, due to the similarity among tested sgRNAs, we confirmed that each experiment was associated with at most one sgRNA. As a result, we generated 20 new GUIDE-seq experiments with 646 OTS, including 147 sites containing bulges across the hg38 reference genome (Table 1). Our GitHub repository, CRISPR-Bulge, contains the processed data and the DNA barcodes in their correct association.

During our analysis, we modified the GUIDE-seq pipeline to support parallel computing and speed up the process. The modified version is available at github. com/ofiryaish/guideseq_parallelized.

#### Independent GUIDE-seq dataset

To gauge the prediction performance on independent GUIDE-seq datasets, we analyzed GUIDE-seq experiments which were curated by crisprSQL (32): Tsai 2015 (15) (conducted on HEK293 and U2OS cells), and Kleinstiver 2016 (33), Chen 2017 (34), and Listgarten 2018 (24) (conducted on U2OS cells). For consistency with our GUIDE-seq data processing protocol, and since some of those datasets lacked OTS with bulges due to the outdated pipeline, we applied the updated pipeline to their raw data. DNA barcodes were not publicly available for Tsai 2015 and Kleinstiver 2016. Upon request, the generators of Tsai 2015 provided us with the barcodes, while the generators of Kleinstiver did not.

Overall, for the independent GUIDE-seq dataset, we processed 39 GUIDE-seq experiments with a total of 711 OTS, including 142 sites containing bulges across the hg38 reference genome (Table 1). The processed data and DNA barcodes are available in our CRISPR-Bulge GitHub repository.

#### Generating active and inactive OTS for machine-learning models

An essential step in developing an effective off-target predictor is constructing datasets that include active and inactive OTS (i.e., sites obtained experimentally and potential sites based on sequence similarity that were not cleaved) (35). To obtain all potential OTS, we employed SWOffinder, a novel method we developed for searching CRISPR OTS with bulges (36). Compared to previous search methods such as Cas-OFFinder (37), which were utilized in CRISPR-Net and CRISPR-IP (26; 29), SWOffinder guarantees the detection of all potential OTS of Cas9 RNA-guided endonucleases, both with and without bulges, given a specific sgRNA and a reference genome. Based on the number of mismatches and bulges seen in the experimental data and common thresholds used in previous studies (26; 29), we configured the search parameters to identify all potential OTS with up to 1 DNA or RNA bulge and 4 mismatches or with up to 6 mismatches in the absence of bulges compared to the given sgRNA.

To generate the inactive OTS, we filtered out experimentally identified OTS from the pool of potential OTS obtained by SWOffinder across the hg38 reference genome. Specifically, since each off-target site can be uniquely identified by the quartet (sgRNA, chromosome, strand, and genome end position), we removed potential OTS from the list if their quartet identifier matched those of the experimentally identified OTS. Additionally, for each sgRNA, we excluded potential OTS found within a genome window surrounding the experimentally identified sites for that particular sgRNA. This approach is not only based on the assumption that neighboring sites may be considered the same, especially when considering bulges, but also takes into account the fact that current state-of-the-art experimental techniques, such as CHANGE-seq and GUIDE-seq (15; 30), cannot precisely determine the exact position of an off-target site within a sequencing read. Similar to the GUIDE-seq pipeline, we defined the genome window as the identified off-target site plus an additional 25 flanking positions. We consider all potential OTS that satisfy the search thresholds in a window as inactive OTS. When multiple optimal alignments exist for the same target site, SWOffinder outputs the alignment with fewer bulges. Table 1 reports the total number of inactive OTS for the CHANGE-seq and GUIDE-seq datasets.

We defined experimentally obtained OTS as the active sets. In the CHANGE-seq study and our previous work (35), experimentally identified OTS cleaved *in vitro* with read counts below 100 were filtered out as noisy measurements. This filtering criterion resulted in a reduced size of the CHANGE-seq active set to 70, 425 OTS. Thus, we re-examined this filtering criterion. We did not reprocess the active OTS of the CHANGE-seq dataset.

#### The refined TrueOT benchmark

Experimentally validated OTS are generated using HTS on PCR amplicons techniques, such as rhAmpSeq (38), which are considered the gold standard for validating OTS (39). Inspired by the TrueOT dataset (39; 40), a comprehensive collection of 11 different studies designed specifically for evaluating experimentally validated OTS in living cells, we chose for our refined TrueOT benchmark 4 studies (41; 42; 43; 30) that satisfy two criteria: (i) the amplicon-based HTS assay was generated following a genome-wide *in cellula* unbiased assay (e.g., GUIDE-seq), which allowed us to include inactive OTS from across the genome. (ii) the assays were not used to train the models participating in our prediction performance evaluation. Similar to previous studies, we labeled OTS with an editing rate greater than 0.1% as active OTS, and defined inactive OTS using the same process as before to consider the entire genome (Sub-section A). For consistency with our GUIDE-seq data generation pipeline, we also applied realignment of the active OTS. Since not all OTS obtained in the GUIDE-seq experiments of the CHANGE-seq study were experimentally validated (30), we excluded such OTS from both the inactive and active OTS in the refined TrueOT dataset. Overall, the refined TrueOT benchmark contains 10 GUIDE-seq experiments with a total of 100 active OTS and 354, 252 inactive OTS (Table 1).

### B. Our deep neural networks to predict OTS

#### Sequence encoding

In our deep neural networks, the prediction of off-target activity is based on aligned sgRNA and off-target sequences, which include the PAM. To accommodate aligned sequences with bulges, we extend the one-hot-encoding approach utilized in our previous study (35), which could only handle mismatches, to handle bulges. Given an aligned sgRNA sequence *S*_*t*_ and an aligned off-target sequence *S*_*o*_, we encode the pair of aligned nucleotides or gaps (represented by -) at position *i* in the aligned sequences into a 5 *×* 5 matrix *C*_*i*_(*j*_*t*_, *j*_*o*_) using the following scheme:

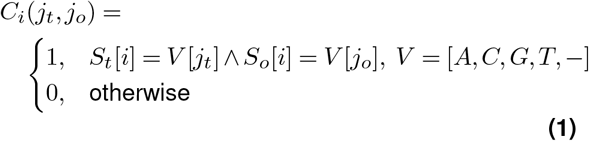

where 1 ≤ *j*_*t*_, *j*_*o*_ ≤ 5 represent the four possible nucleotides and a gap. Prior to the encoding process, we adjust the N in the sgRNA PAM sequence to match the corresponding nucleotide in the off-target site.

To generate one-hot-encoding sequence features for a pair of aligned sequences, each matrix representing the aligned pair at a specific position is flattened. Subsequently, all of these vectors are stacked to form a one-hot-encoding matrix. As the alignment length can be either 23 or 24, shorter alignments are left-padded with a pair of gaps. This results in a fixed-sized one-hot-encoding matrix of dimensions 25 *×* 24, which is then used as input to our models (Figure 1A).

**Figure 1.**
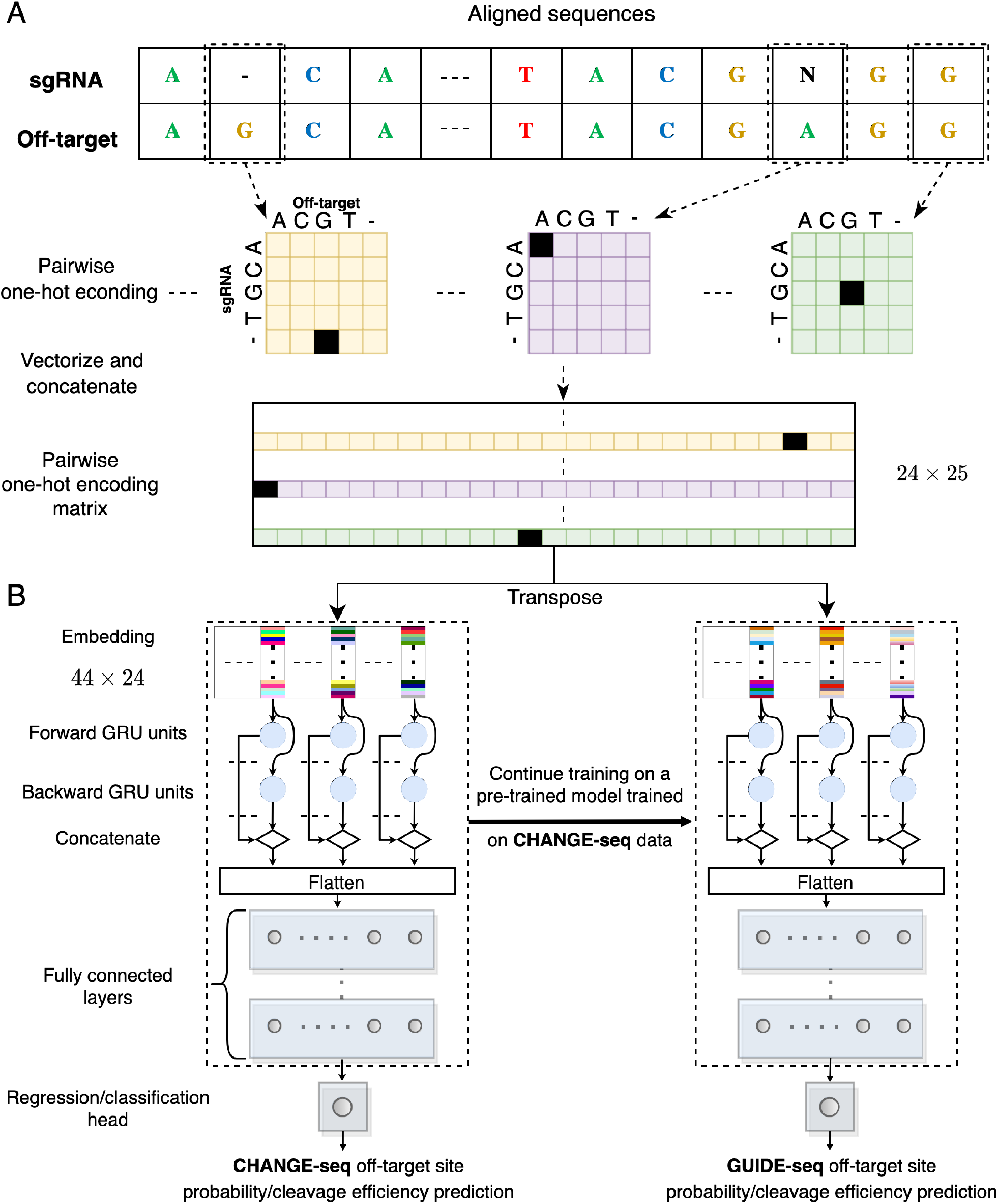
Predictive model overview. **(A-B)** Illustration of the GRU-Emb architecture and its training process. Given an aligned pair of sgRNA and off-target site sequences, the sequences are processed using pairwise one-hot encoding, and then the flattened positional vectors are stacked (A). Subsequently, the fixed-sized one-hot-encoding matrix is provided as input to the GRU-Emb classifier/regressor to predict the off-target site probability/cleavage efficiency (B). A transfer-learning approach utilizes the large-scale CHANGE-seq dataset to train a dedicated model for the GUIDE-seq data.

#### Classification and regression models

We developed two types of predictive models. The first, a classification model, aims to predict the cleavage probability of an off-target site. The second, a regression model, aims to predict the cleavage efficiency of an off-target site. In the case of a classification model, we defined the positive and negative sets as the off-target active and inactive sets. For the regression model labels, we employed a data transformation technique where the read counts, denoted as *x*, were log-transformed using the formula *log*(*x* + 1) (35). We assigned inactive OTS a read count of 0 to reflect their lack of cleavage activity.

#### Deep neural network architectures

We trained three deep-neural-network architectures (inspired by pi-CRISPR (44) and DeepHF (45) architectures) for off-target classification and regression tasks, leveraging multilayer-perceptron (MLP), gated-recurrent-unit (GRU), and embedding techniques. The first architecture, which we denote by *MLP-Emb*, begins with an embedding layer of size 44, which transforms each 25-long one-hot-encoded aligned nucleotide pair into a 44-long real vector representation. The embedded sequences are then processed by a flattening layer. Subsequently, the flattened features pass through two consecutive dense layers with 128 and 64 neurons with ReLU activation function. The last layer is a single neuron with a sigmoid or linear activation for the classification and regression tasks, respectively. The second architecture, which we denote *GRU-Emb*, closely resembles the MLP-Emb architecture. However, the embedded input sequences are fed into a GRU layer before being processed by the dense layers (Figure 1B). We set the number of GRU units to 64. The third architecture, which we denote *GRU*, is similar to GRU-Emb but without the embedding layer. Hyper-parameters values of the models, such as the embedding size, were similarly inspired by piCRISPR and DeepHF. We denote classification and regression models by the suffix Class or Reg, respectively.

#### Model training and transfer learning to GUIDE-seq

To ensure consistency and prevent mixing datasets with different characteristics, we trained separate models for the CHANGE-seq and GUIDE-seq datasets. In addition to training a separate model solely on the GUIDE-seq data, we employed a transfer-learning approach to utilize the large-scale CHANGE-seq dataset to train a dedicated model for the GUIDE-seq data. In this technique, we first trained our model on the CHANGE-seq data and then continued the training on the GUIDE-seq data. We denote models trained solely on CHANGE-seq or GUIDE-seq by the prefix CH or GU, respectively, and the transfer-learning model by the prefix TL.

To improve the prediction performance and robustness of our models, we applied the random ensemble initialization (46). We trained the same architecture with 5 different random initializations and output the average over their predictions in test time. In our leave-k-sgRNA-out cross-validation evaluations, we did not use the ensemble technique to reduce training runtime, and since the k-fold approach already reduced the variance and made the results more robust (47).

We trained the models using the Adam optimizer with a learning rate of 0.005. We used a batch size of 512 with a maximum number of training epochs set to 10. We utilized the binary cross-entropy and mean-squared-error loss functions for the classification and regression tasks, respectively. During each model training, we used a validation set comprising 10% of the training data to select the optimal epoch based on the validation loss. Considering the highly imbalanced nature of both the GUIDE-seq and CHANGE-seq datasets, we investigated the effectiveness of class weighting in neural network models, a technique we previously used in an XGBoost model (35). For each active or inactive set, we defined its weight to be proportional to the size of the other set.

### C. Prediction performance evaluations

#### Dataset partitioning to training and test sets

In all of our evaluations, no sgRNA is found in both the training and test sets, even across experimental techniques. For the CHANGE-seq and GUIDE-seq datasets of the CHANGE-seq study, we used a shared partition when evaluating the prediction performance of the models to avoid sgRNA overlap. We first partitioned the dataset of 58 GUIDE-seq sgRNAs into 10 sets, each containing samples from 5 or 6 sgRNAs. As 28 of the GUIDE-seq sgRNAs have active OTS with bulges, we used a greedy approach to maintain that each part in the partition has samples of 2 or 3 sgRNAs and the number of active OTS with bulges in the sets is nearly balanced (see our CRISPR-Bulge GitHub repository for the data partition). For the partition over the CHANGE-seq dataset, we augmented the partition over the GUIDE-seq dataset by randomly adding sgRNAs from the remaining 52 sgRNAs to each part to a size of 11.

When evaluating the TL and CH models on the new GUIDE-seq dataset, we excluded any CHANGE-seq data of the 20 sgRNAs shared with the new GUIDE-seq dataset. For the prediction performance evaluation over the independent GUIDE-seq and the refined TrueOT datasets, we used the same CH and TL models where the training set is comprised of the entire CHANGE-seq, GUIDE-seq, and new GUIDE-seq data, except for\ OTS of 6 sgRNAs included in the refined TrueOT as rhAmpSeq experiments.

#### Evaluation metrics

We evaluated prediction performance over multiple sgRNAs. On the CHANGE-seq and GUIDE-seq datasets, we used our 10-fold sgRNA-based partition for evaluation. During each iteration of a cross-validation process, we used one fold as the test set for evaluation, and the remaining 9 folds as the training set. We performed the evaluation of the new GUIDE-seq, independent GUIDE-seq, and the refined TrueOT datasets on the entire datasets.

We evaluated the performance of the models in the classification task using the area under the precision-recall curve (AUPR, calculated as average precision), which is appropriate for imbalanced test cases and recommended for evaluation of OTS prediction performance (48), and area under the ROC curve (AUC), which is a common binary-classification evaluation metric. In addition, we defined a new evaluation by assessing performance on OTS with bulges only. Unlike previous studies that focused on the overall dataset performance, we sought to provide increased resolution to the evaluation by focusing on the more challenging task of OTS with bulges.

#### Comparison to current methods for off-target prediction

For a baseline comparison, we implemented a modified version of the XGBoost model used in previous studies (30; 35) to enable the prediction of OTS with bulges. The sequence encoding is as the encoding for our deep neural network models, but with an additional flattening due to the limitation of XG-Boost in processing two-dimensional data. To conduct more advanced comparisons (i.e., on the TrueOT dataset), we compared our models to the state-of-the-art methods that can handle OTS with bulges, COSMID (49), CRISTA (21), CRISPR-Net (26) and CRISPR-IP (29). We tried running CRISPR-HW (50), CRISPR-BERT (51), and CRISPR-M (52), but the software, as downloaded from the GitHub repositories, failed to run due to various reasons: CRISPR-HW and CRISPR-M did not provide a trained model, and CRISPR-BERT raised executions errors when performing the prediction.

### D. Model interpretation

#### Visualizing the positional effect on off-target activity

To visualize the positional effect of mismatches and bulges on off-target activity, we randomized 100 sgRNAs and, for each one, generated all possible OTS with a single mismatch or a DNA/RNA bulge, and all double-mismatched OTS. To assess the positional effect, we averaged the predictions across the 100 sgRNAs for each type of edit (mismatch, DNA/RNA bulge, and double mismatch) and plotted the average prediction and standard deviation as a function of the edit position. For the single edit, we also visualized the effect of a specific nucleotide mismatch or DNA/RNA bulge. In addition, similar to (53), we examined the *epistasis-like combinatorial effect* of two mismatches. We compared the prediction of our models to a simplified theoretical model where the prediction is the multiplication of individual mismatch predictions (i.e., assumes the combinatorial effect of two or more mismatches is marginally independent). We performed the analyses on both the CHANGE-seq and GUIDE-seq models to assess the consistency between the principles learned from *in vitro* and *in cellula* data.

#### Visualizing the learned representation

To understand how the embedding models (i.e., MLP-Emb and GRU-Emb) capture the differences between the different types of OTS, we visualized the representation learned by the models. We randomly selected 20 sgRNAs from the CHANGE-seq dataset and for each sgRNA, randomly generated 1, 000 and 2, 000 OTS *in silico* with up to 4 edits without bulges and with 1 DNA/RNA bulge, respectively. We then employed T-distributed stochastic neighbor embedding (t-SNE) to project the high-dimensional embedded representations into a two-dimensional map (54). We average the embedding vectors of the models in the ensemble. To reduce the dependency on the different sgRNAs, we subtract the embedding of the on-target site from the embeddings of its OTS. For comparison, we repeated this visualization analysis using the one-hot-encoding sequence features.

## Results

### E. A new GUIDE-seq dataset including hundreds of CRISPR off-target sites with bulges

We generated the largest dataset to date of CRISPR OTS with bulges (Table 1). We retrieved data of 97 previously published GUIDE-seq experiments from various sources, extracted novel OTS with bulges, and generated 20 new GUIDE-seq experiments (Subsection A). As a result, we expanded the set of GUIDE-seq datasets to cover 117 sgRNAs, with a combined total of 2, 877 identified OTS, including 591 OTS with bulges. To the best of our knowledge, this is the most comprehensive set of GUIDE-seq datasets containing OTS with bulges to date.

### F. Improved prediction of CRISPR off-target sites with bulges

We first explored the effect of removing the read-count threshold, using class weighting in the training process, and input encoding of our models. We evaluated the models by a leave-11-sgRNAs-out cross-validation evaluation on the CHANGE-seq dataset. To perform an objective comparison when examining the removal of the read-count threshold and for consistency with previous studies, we evaluated the performance on the subset of OTS with a read-count threshold of 100.

The models exhibited improved performance when the read-count threshold was removed (Figures 2A-B). For instance, the CH-GRU-Reg model without class weighting and thresholding achieved an average AUPR of 0.636, whereas the comparable model with thresholding achieved an average AUPR of 0.594 (p-value = 1.95 *×* 10^−3^ via Wilcoxon signed-rank test). Thus, we proceeded with removing the read-count threshold in all model training procedures. The full results on each test fold are reported in Supplementary Table S1.

**Figure 2.**
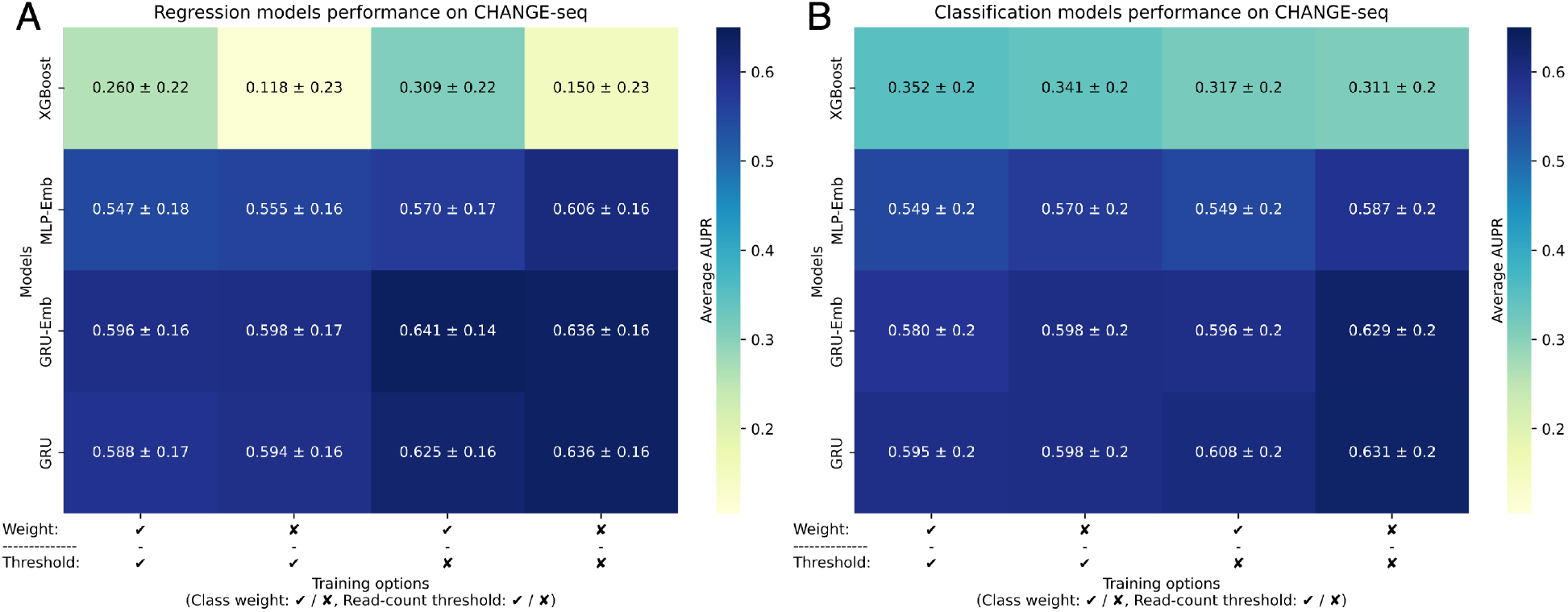
Prediction performance evaluations on *in vitro* data. **(A-B)** A comparison of the CH-XGBoost, CH-MLP-Emb, CH-GRU, CH-GRU-Emb regression (A) and classification (B) models on the CHANGE-seq data examining the effect of using class weighting and read-count threshold. We gauged the prediction performance of the models by average AUPR over the leave-11-sgRNAs-out cross-validation folds.

We observed opposite effects of class weighting on the XGBoost and neural network models. Consistent with our previous study (35), the classification and regression XGBoost models benefit from class weighting in all model variations. However, when examining the neural network models, we found no such improvement and even a decrease in the average AUPR for most models. For example, the CH-GRU-Class model without class weighting and thresholding achieved an average AUPR of 0.631, while the same model with class weighting achieved an average AUPR of 0.598 (p-value = 0.049). Thus, in subsequent model evaluations, we do not apply class weighting in training. The full results on each test fold are reported in Supplementary Table S1.

As for the input encoding, we compared our encoding of the aligned sgRNA and OTS to the encoding of CRISPR-Net (26). Our evaluation showed that in our training settings (Supplementary Figure S1), the CRISPR-Net encoding with the embedding models failed to predict well (AUPRs *<* 0.1), and also resulted in reduced prediction performance compared to our encoding in other model types (AUPRs ≤ 0.56).

### G. Unique evaluation of CRISPR off-target sites with bulges

We uniquely compared the prediction performance, in the leave-11-sgRNAs-out cross-validation evaluation over the CHANGE-seq dataset, of the classification and regression models on a subset of OTS consisting exclusively of OTS with bulges and on the entire dataset. We focused specifically on the CH-GRU and CH-GRU-Emb models, as the results indicated their superiority over the CH-XGBoost and CH-MLP-Emb models (Figure 2). The computed statistical significance is based on 10 AUPRs of the cross-validation evaluation.

When evaluating the performance on the entire CHANGE-seq dataset and its only-bulges subset (Supplementary Figure S2), the regression models consistently achieved slightly higher average AUPR values than their corresponding classification models, albeit without statistical significance. For example, on the only-bulges subset evaluation, the CH-GRU-Emb-Reg and CH-GRU-Reg achieved an AUPR of 0.498 and 0.492, respectively, compared to the CH-GRU-Emb-Class and CH-GRU-Class models, which achieved an AUPR of 0.491 and 0.467, respectively. Based on these results, we continued with both the regression and classification models for further analysis. The full results on each test fold are reported in Supplementary Table S2.

### H. Improved prediction of CRISPR off-target sites with bulges *in cellula* through transfer learning

Following the training on the CHANGE-seq dataset (i.e., *in vitro* data), we fine-tuned the models to *in cellula* data by continuing the training on the GUIDE-seq data. To assess the effectiveness of our transfer learning, we compared its performance with models trained solely on GUIDE-seq or CHANGE-seq data. The GUIDE-seq data we used for training and testing the models in this evaluation is the subset of 58 sgRNAs that were originally published as part of the CHANGE-seq study.

In both evaluations on the entire and only-bulges GUIDE-seq datasets, we observed that the transfer-learning approach outperformed the other models (Figures 3A-B). For instance, in the entire GUIDE-seq dataset evaluation (Figure 3A), the best performing model was the TL-GRU-Reg, which achieved an average AUPR of 0.432 compared to 0.365 and 0.214 achieved by its corresponding GRU models trained solely on GUIDE-seq and CAHNGE-seq data, respectively (p-values ≤ 1.95 *×* 10^−3^). In the evaluation on the only-bulges dataset (Figure 3B), the TL-GRU-Reg model also showed the best performance with an average AUPR of 0.287, compared to its corresponding GRU models trained solely on GUIDE-seq and CHANGE-seq data, which achieved an average AUPR of 0.156 and 0.165, respectively (p-values *<* 0.049). Overall, all transfer-leaning models performed similarly and outperformed the models trained solely on GUIDE-seq or CHANGE-seq data. The full results on each test fold are reported in Supplementary Table S3.

**Figure 3.**
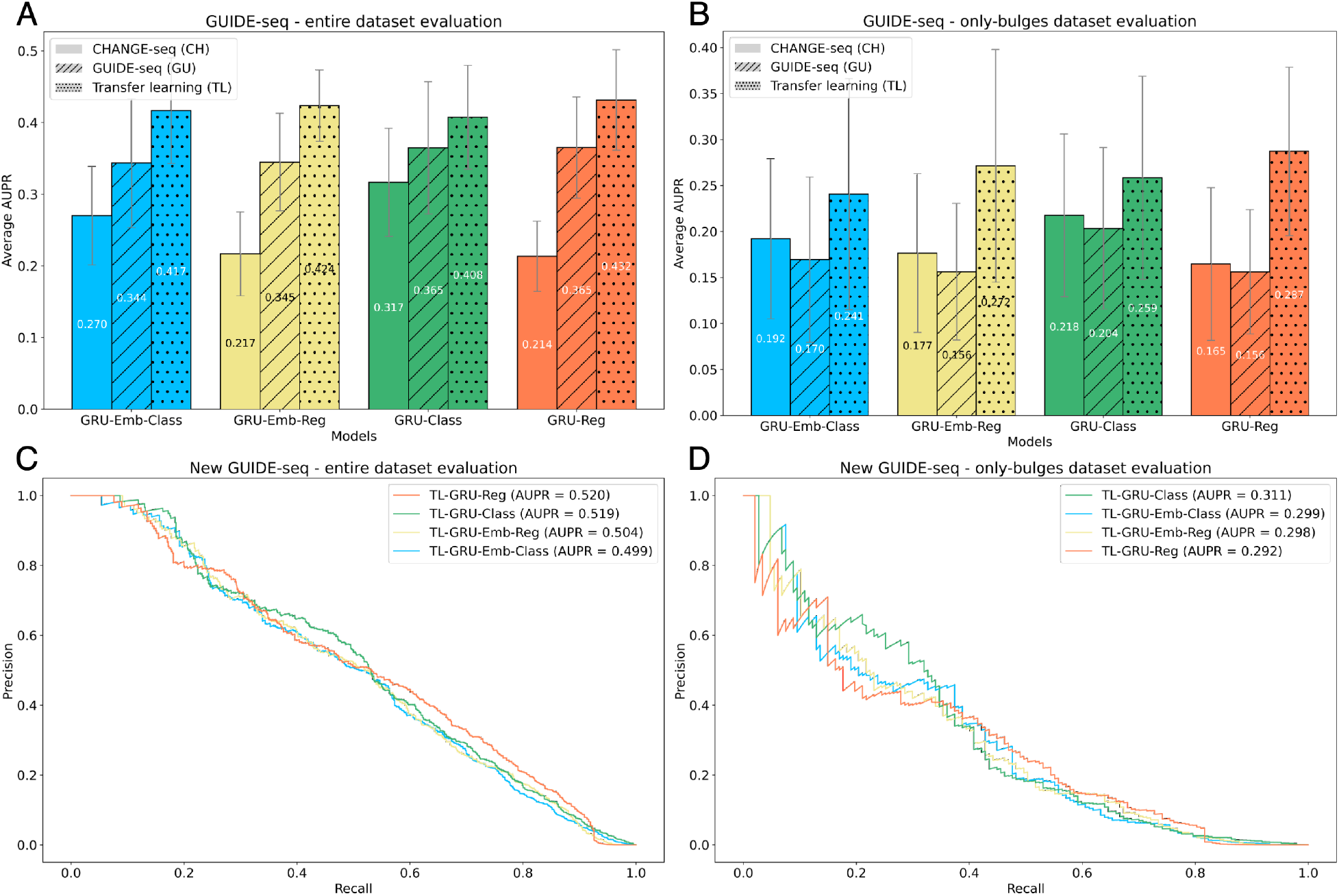
Prediction performance evaluations on *in cellula* data. **(A-B)** Evaluation on the entire GUIDE-seq dataset that was published in the CHANGE-seq study (A) and its only-bulges subset (B). GRU-Reg, GRU-Emb-Reg, GRU-Class, and GRU-Emb-Class models trained on different data are examined: trained only on CHANGE-seq (CH), trained only on GUIDE-seq (GU), and trained on CHANGE-seq and then fine-tuned on GUIDE-seq (TL). **(C-D)** Evaluation of the TL-GRU-Reg, TL-GRU-Emb-Reg, TL-GRU-Class, and TL-GRU-Emb-Class models on our newly generated 20 GUIDE-seq experiments (C) and their only-bulges subset (D).

### I. Evaluation of our newly generated GUIDE-seq dataset

To validate the quality of our 20 newly generated GUIDE-seq experiments, we tested the performance of our transfer-learning models on them. We trained the pre-trained models on the entire CAHGNE-seq dataset, excluding the sgRNAs of the new 20 GUIDE-seq datasets. Then, we continued the training on the subset of 58 GUIDE-seq sgRNAs that were originally published in the CHANGE-seq study.

Overall, the prediction performance on the 20 new GUIDE-seq experiments was on par with the performance achieved on the 58 original GUIDE-seq experiments (Figures 3C-D), indicating the high quality of the new dataset. On the entire new GUIDE-seq dataset (Figure 3C), the TL-GRU-Reg and TL-GRU-Class achieved an AUPR of 0.520 and 0.519, respectively, slightly outperforming the TL-GRU-Reg and TL-GRU-Class models, which achieved an AUPR of 0.504 and 0.499, respectively. Similarly, all the models resulted in comparable performance in the only-bulges evaluation. However, the TL-GRU-Class achieved better prediction performance with an AUPR of 0.311, whereas the other models resulted in similar AUPRs of almost 0.3 (Figure 3D).

### J. Benchmarking methods on an independent GUIDE-seq dataset

To evaluate the prediction performance of the models on independent GUIDE-seq experiments, that were not conducted on T cells and not included in the CHANGE-seq study, we compared our transfer-learning models trained on the entire CHANGE-seq and GUIDE-seq datasets (with the exclusion of a few sgRNAs to avoid shared sgRNAs in train and test sets in our evaluation on the refined TrueOT dataset which shares the same trained models with this evaluation, Subsection C) to state-of-the-art prediction methods that can handle bulges, COSMID, CRISTA, CRISPR-Net, and CRISPR-IP (49; 21; 26; 29) on our independent GUIDE-seq dataset comprised of 3 GUIDE-seq datasets: Tsai 2015, Chen 2017, and Listgarten 2018 (Subsection A).

CRISPR-Net and CRISPR-IP were trained on the Tsai 2015 and Listgarten 2018 GUIDE-seq datasets, and thus had an advantage over the other methods.

Overall, our models outperformed all tested methods by a large margin in the entire dataset evaluation and in both the only-bulges and only-mismatches subsets (Figures 4A-C). The TL-GRU-Emb-Class and TL-GRU-Class achieved the highest AUPRs of 0.466 and 0.443 in the entire dataset evaluation, respectively, outperforming (among all other methods) our TL-GRU-Reg and TL-GRU-Emb-Reg, which achieved an AUPR of 0.342 and 0.306, respectively. Similar prediction performance trends were obtained separately on each GUIDE-seq dataset (Supplementary Figure S3) and when evaluating the performance using AUROC (Supplementary Figure S4). Moreover, our transfer-learning approach outperformed matching models trained solely on the CHANGE-seq dataset or on the dataset used in the CRISPR-Net method (Supplementary Figure S5).

**Figure 4.**
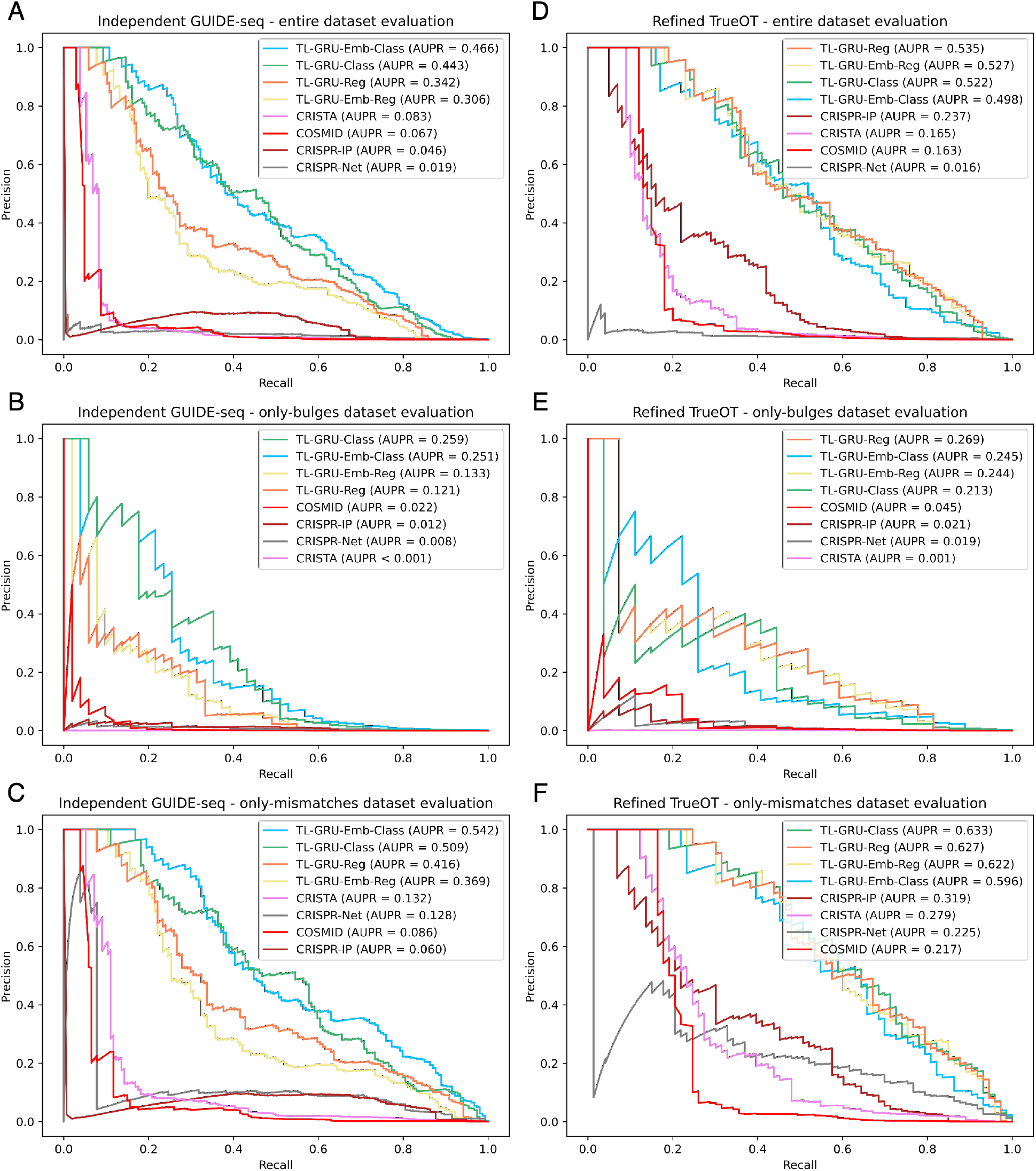
Prediction performance evaluations on independent GUIDE-seq and experimentally validated datasets. **(A-C)** Comparison of our models TL-GRU-Reg, TL-GRU-Emb-Reg, TL-GRU-Class, and TL-GRU-Emb-Class with state-of-the-art methods COSMID, CRISTA, CRISPR-Net, and CRISPR-IP on our independent GUIDE-seq benchmark dataset (A), its only-bulges subset (B), and its only-mismatches subset (C). **(D-F)** Comparison of our models with state-of-the-art methods on our refined TrueOT dataset (D), its only-bulges subset (E), and its only-mismatches subset (F).

### K. Benchmarking methods on an experimentally validated off-target sites dataset

In our final prediction performance evaluation, we evaluated the prediction performance of our models trained on the entire CHANGE-seq and GUIDE-seq datasets through transfer learning (with the exclusion of a few sgRNAs to avoid shared sgRNAs in train and test sets in our evaluation, Subsection C) on our refined TrueOT dataset. This dataset contains experimentally validated OTS generated using HTS on PCR-amplicons techniques, such as rhAmpSeq (38), which are considered the gold standard for validating OTS (39). Our refined TrueOT dataset contains 354, 352 potential OTS, 100 of which are experimentally validated, including 27 with bulges (Subsection A).

Our TL-GRU-based models outperformed COSMID, CRISTA, CRISPR-Net, and CRISPR-IP (49; 21; 26; 29), where the most superior advantage was achieved in the evaluation on the subset containing only OTS with bulges (Figures 4D-F). On the entire refined TrueOT dataset (Figure 4D), the TL-GRU-Reg, TL-GRU-Emb-Reg, TL-GRU-Class, and TL-GRU-Emb-Class models achieved similar performance with an AUPR of 0.535, 0.527, 0.522, and 0.498, respectively, outperforming CRISPR-IP, CRISTA, COSMID, and CRISPR-Net models with an AUPR ≤ 0.237. Similarly, in the only-bulges evaluation, our TL-GRU-based models achieved an AUPR ≥ 0.213, outperforming the competing methods by a large margin: an AUPR ≤ 0.045 (Figure 4E). Our models achieved improved prediction performance even in the only-mismatches evaluation (Figure 4F). The same trends were observed when evaluating the prediction performance by the AUROC metric (Supplementary Figure S6). The TL-GRU-based models also outperformed matching models trained solely on the CHANGE-seq dataset or on the dataset used in the CRISPR-Net method (Supplementary Figure S7).

### L. Analyzing positional effect on off-target activity

We analyzed the positional effect of mismatches and bulges on off-target activity learned by our GRU-Emb-Class model trained on *in vitro* or *in cellula* data. We chose the GRU-Emb-Class model since the classification models outperformed the regression models overall and since we wanted to inspect what the embedding layer has learned. We examined the positional effect of a single edit compared to the sgRNA (Figure 5A-F). As expected, the models for *in vitro* and *in cellula* data learned that edits near the PAM are less likely in active OTS. As an exception, the *in cellula* model learned that edits in positions 15-17 are less tolerable than edits in positions 13-14 and 18-19 (Figures 5B, 5D, and 5F). In addition, the *in cellula* model showed sensitivity around position 5, especially for bulges. These patterns were also learned by the GRU-Emb-Reg models (Supplementary Figure S8). These position-dependent patterns agree with findings observed in previous high-throughput synthetic assays (55; 56). The *in vitro* model also captured similar patterns for positions around position 16, albeit with reduced intensity (Figures 5A, 5C, and 5E). Another dissimilarity between the *in vitro* and *in cellula* models is the higher sensitivity of the *in cellula* model to edits near the PAM. When examining the effect of a specific mismatch or RNA/DNA bulge, we found that around position 16, there is an increased intolerance for mismatches when the sgRNA nucleotide is C or when the OTS nucleotide is G or T (Supplementary Figure S9).

**Figure 5.**
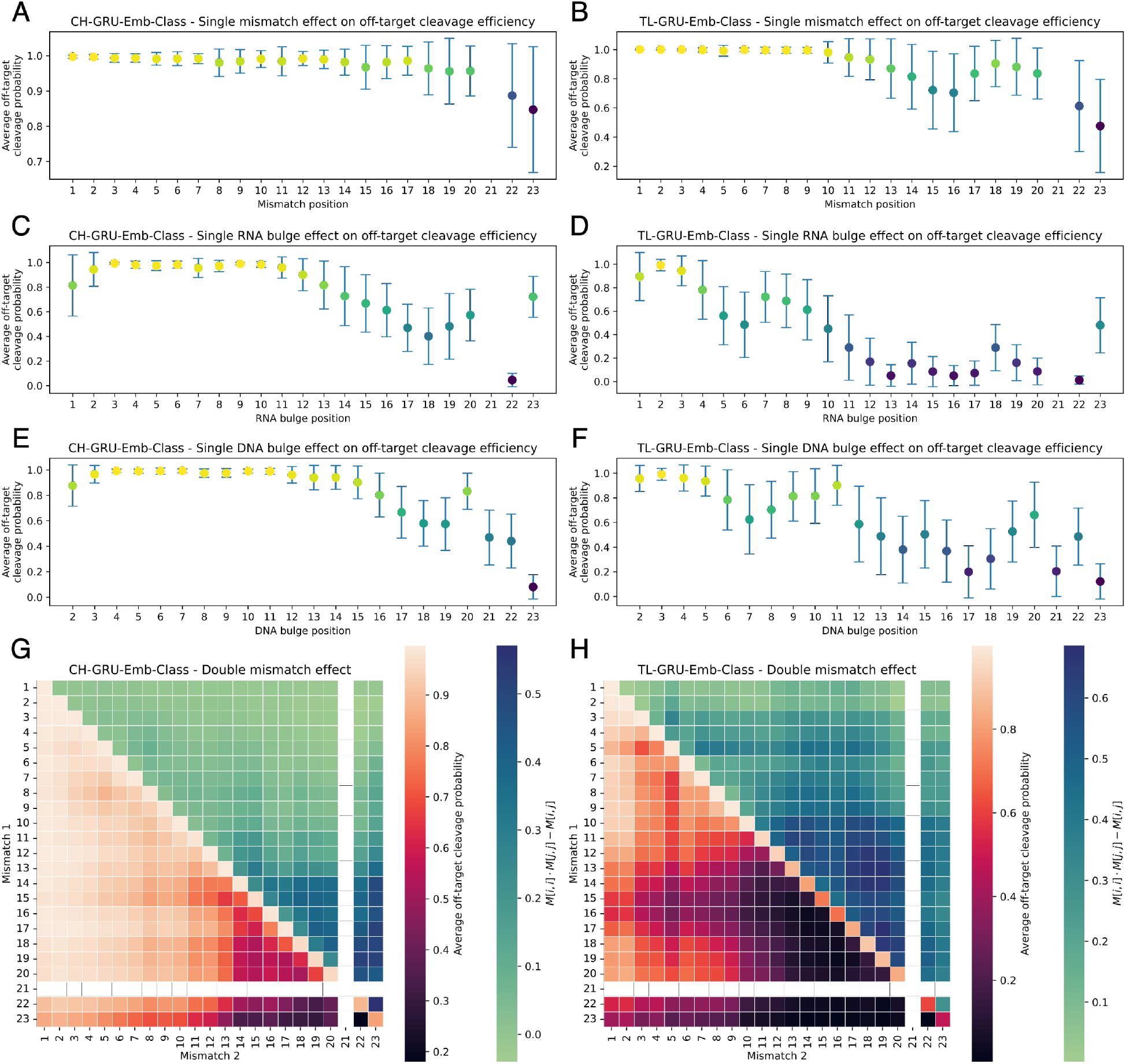
Positional effect of mismatches and bulges on off-target activity learned by our GRU-Emb-Class model. **A-F** Positional effect of a single mismatch (A-B), RNA bulge (C-D), or DNA bulge (E-F) compared to the sgRNA, predicted by the CH-GRU-Emb-Class and TL-GRU-Emb-Class models for *in vitro* (A, C, and E) and *in cellula* (B, D, and F) data, respectively. **G-H** Positional effect of two mismatches (lower triangular) and the epistasis-like combinatorial effect of two mismatches (upper triangular) compared to the sgRNA predicted by the CH-GRU-Emb-Class (G) and TL-GRU-Emb-Class (H) models.

Furthermore, we analyzed the positional effect of two mismatches compared to the sgRNA (Figure 5G-H). Overall, the general patterns learned in the single mismatch case agree with the double-mismatch trends. However, in contrast to the single mismatch scenario, where a similar trend was learned by the GRU-Emb-Class models for both *in vitro* or *in cellula*, the *in cellula* model exhibited increased intolerance to more than one mismatch, even at positions distant from the PAM. Similar trends were observed when performing the same analysis using the GRU-Emb-Reg model (Supplementary Figure S8). Regarding the epistasis-like combinatorial effect of two mismatches (53), the *in cellula* and *in vitro* models learned that this effect is stronger between mismatches at positions closer to the PAM compared to positions distant from the PAM (Figure 5G-H).

### M. Visualization of the learned embedded representation of off-target sites with bulges

To understand what the network has learned, we visualized the output of the embedding layer for various OTS of randomly selected 20 sgRNAs from the CHANGE-seq dataset in a t-SNE plot. Figure 6 visualizes the learned embedded representation by the TL-GRU-Emb-class model projected into a two-dimensional map using t-SNE following subtraction of the sgRNA on-target representation. Without this subtraction, t-SNE clusters OTS from different sgRNAs as unique clusters (Supplementary Figure S10A). Compared to an equivalent visualization based on one-hot-encoding sequence features (Supplementary Figure S10B), the embedding-based visualization effectively showcases a clear separation of OTS based on the presence or absence of bulges. Moreover, it distinctly illustrates the separation between OTS, featuring different types of bulges.

**Figure 6.**
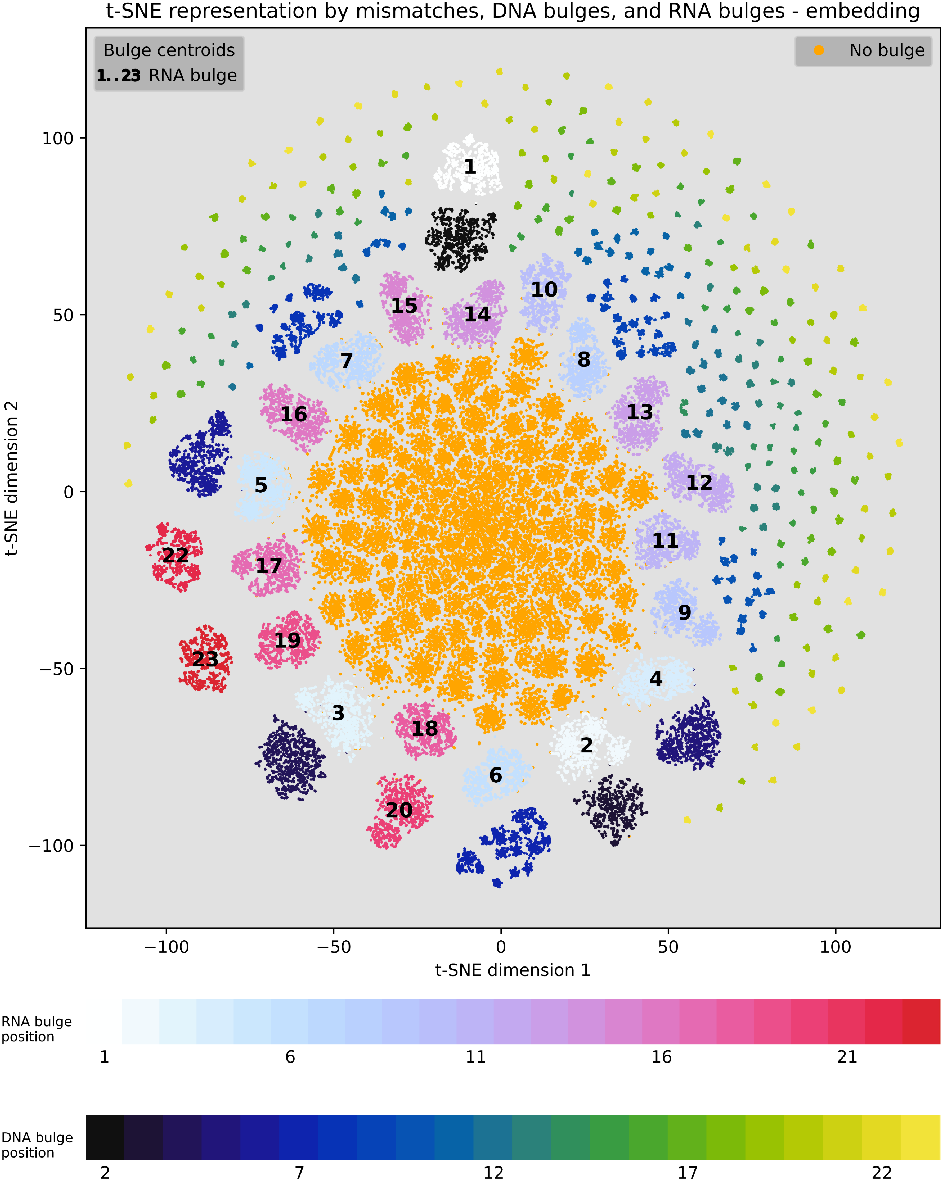
Visualization of the learned embedded representation of off-target sites. Learned embedded representation by the TL-GRU-Emb-Class model projected onto a two-dimensional map using t-SNE. Comparing OTS with mismatches to OTS with bulges according to their bulge type and position. The centroid of OTS with the RNA bulge position is illustrated in each map by the position of the label, while the centroid of DNA bulge positions are dismissed for visual clarity.

The observed separation in the learned embedded space can provide insights into model prediction preferences assuming that proximity in the learned embedded space implies similar predictions by the model. While the model treats OTS without bulges similarly, as their cluster is very dense, it treats OTS with bulges differently according to bulge type and position. Generally, OTS with bulges positioned farther from the PAM exhibit closer proximity to the cluster of OTS without bulges. Surprisingly, clusters of OTS with an RNA bulge are in a radius from the center of the plot, which depends on their distance from the PAM. This may represent an artifact rather than real biology as it is also present in the plot based on one-hot encoding (Supplementary Figure S10B). In conclusion, OTS with bulges close to the PAM are less likely to be active, reaffirming previous findings obtained by high-throughput synthetic assays (55; 57).

## Discussion

In this study, we made several contributions to the CRISPR/Cas9 off-target research field. First, to the best of our knowledge, we generated the largest dataset of OTS with bulges to date. We reprocessed previously published and new in cellula data, which resulted in hundreds of sites with bulges, an order of magnitude more than in previous datasets. Second, using the newly generated datasets, we trained deep neural networks to predict off-target activity by utilizing embedding and recurrent neural networks with transfer-learning techniques to benefit from both *in vitro* and *in cellula* data. Our networks outperformed state-of-the-art methods by a large margin on independent GUIDE-seq datasets and on an experimentally validated OTS dataset, which is considered the gold standard for validating OTS (39). Third, we defined a new metric to evaluate models for off-target prediction with bulges, focusing on active and inactive OTS with bulges. This allows the close inspection of methods’ performance when considering bulges, which may be obscured when evaluating on all the data. Fourth, we analyzed the positional effect and epistasis-like combinatorial effect of mismatches and bulges on the off-target activity learned by our model, reaffirming previous knowledge and providing new insights. Fifth, thanks to the embedding layer, we developed a new visualization technique that utilizes the learned representation of OTS with bulges and normalizes different sgRNAs, and through the visualization, identified the relevance and importance of bulges as learned by the model.

Our study has several limitations. First, while we made a breakthrough in the scale of GUIDE-seq datasets, and in the scale of OTS with bulges in them, this data is still limited compared to other genomic tasks, where tens of thousands of measurements are available. We tackled this limit by transfer learning from the CHANGE-seq dataset, but more GUIDE-seq data is expected to improve the prediction performance of our models. Second, while we evaluated and compared our models to the state of the art on a comprehensive dataset of experimentally validated OTS, this data still needs to be improved in scale, and additional low-throughput experimental validation of specific OTS is needed for robust benchmarking. Third, a prominent limitation lies in the dependence of our method and data-generation pipelines on the alignment between the sgRNA and the off-target site. This alignment has a critical effect on both the data generation as the pipeline identifies optimal sites in a window, and the machine-learning model as it receives the alignment as input.

Several aspects of our study require further research in the future. First, in addition to the data we used, various experimental assays developed to measure OTS in high-throughput were not used in this study (55; 53; 56). Integrating them, either by data combination or by a meta-predictor that considers the predictions of models trained on each data separately, may improve prediction performance. Second, the efficiency of assays for measuring the genome-wide activity of CRISPR/Cas9 nucleases *in cellula* is limited by chromatin accessibility and other epigenetic factors (31; 58; 44). Therefore, including such information in our models may provide more insights regarding their potential for future improvements. Third, current methods to predict OTS assume an alignment between the sgRNA and the off-target, but multiple alignments may exist. Having a method that receives as input multiple possible alignments, or no alignment at all, may improve prediction performance and shed more light on the interplay between sgRNAs and their OTS. The proposed alignment-free analysis can be further extended to discover the optimal mismatch and bulge thresholds. Fourth, while our main focus was on data generation and our transfer-learning approach, additional work on choosing the optimal model architecture and hyper-parameters may further improve prediction performance.

## Conclusion

We presented a groundbreaking contribution to the CRISPR/Cas9 gene-editing field by introducing the most comprehensive dataset of OTS with bulges. By curating and analyzing a vast collection of experimental data, our study sheds new light on the mechanisms underlying off-target effects induced by CRISPR/Cas9. The inclusion of bulges expands our understanding of the complex interplay between sgRNAs and OTS, providing insights into the factors influencing off-target activity. The availability of our extensive dataset will significantly benefit the scientific community, enabling researchers to develop more accurate prediction models and refine the design of CRISPR/Cas9-based therapies. Ultimately, this research advances the ongoing efforts to enhance the precision, safety, and efficacy of CRISPR/Cas9 technology.

## Author contributions statement

OY conducted all the analyses and developed all the codes used in this study. YO supervised this study. OY and YO wrote the manuscript.

## Acknowledgements

We thank our colleagues at the CRISPR-IL consortium for thoughtful discussions over the off-target prediction problem. Ofir acknowledges the support of the Azrieli Ph.D. fellowship, Nehemia Levtzion Ph.D. fellowship from the Israeli Council for Higher Education, Kretiman Ph.D. fellowship at Ben-Gurion University, and the cloud-compute credit by the Planning and Budgeting Committee through IUCC. We would like to thank the anonymous reviewers for their constructive comments.

## Data Availability

The software and code are publicly available via github.com/OrensteinLab/CRISPR-Bulge. Supplementary Table S4 contains processed data of active OTS for all the datasets: CHANGE-seq, GUIDE-seq, new GUIDE-seq, independent GUIDE-seq (i.e., Tsai 2015, Chen 2017, and Listgarten 2018), and the Refined TrueOT. Complete tables that include inactive OTS are available in the GitHub repository. DNA barcodes, accession codes, and raw data links required for the GUIDE-seq pipeline are available in the GitHub repository. The source code was deposited in Zenodo (DOI: 10.5281/zenodo.10902355).

## Supplemental Data

### N. Supplementary Figures

Supplementary Figures S1-10.

### O. Supplementary Tables

Supplementary Tables S1-4.

## Supplementary Figures

**Figure S1.**
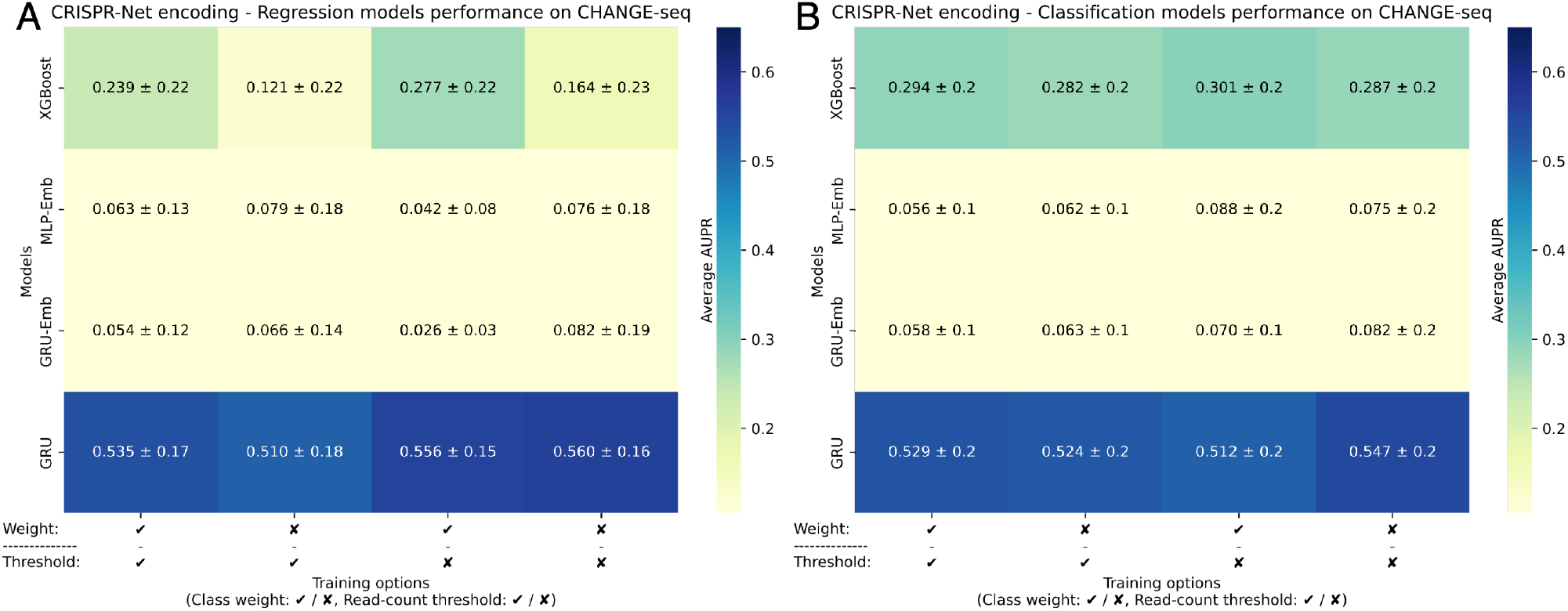
CRISPR-Net encoding prediction performance evaluations on *in vitro* data. **(A-B)** A comparison of the CH-XGBoost, CH-MLP-Emb, CH-GRU, CH-GRU-Emb regression (A) and classification (B) models trained with CRISPR-Net encoding on the CHANGE-seq data examining the effect of using class weighting and read-count threshold. We gauged the prediction performance of the models by average AUPR over the leave-11-sgRNAs-out cross-validation folds.

**Figure S2.**
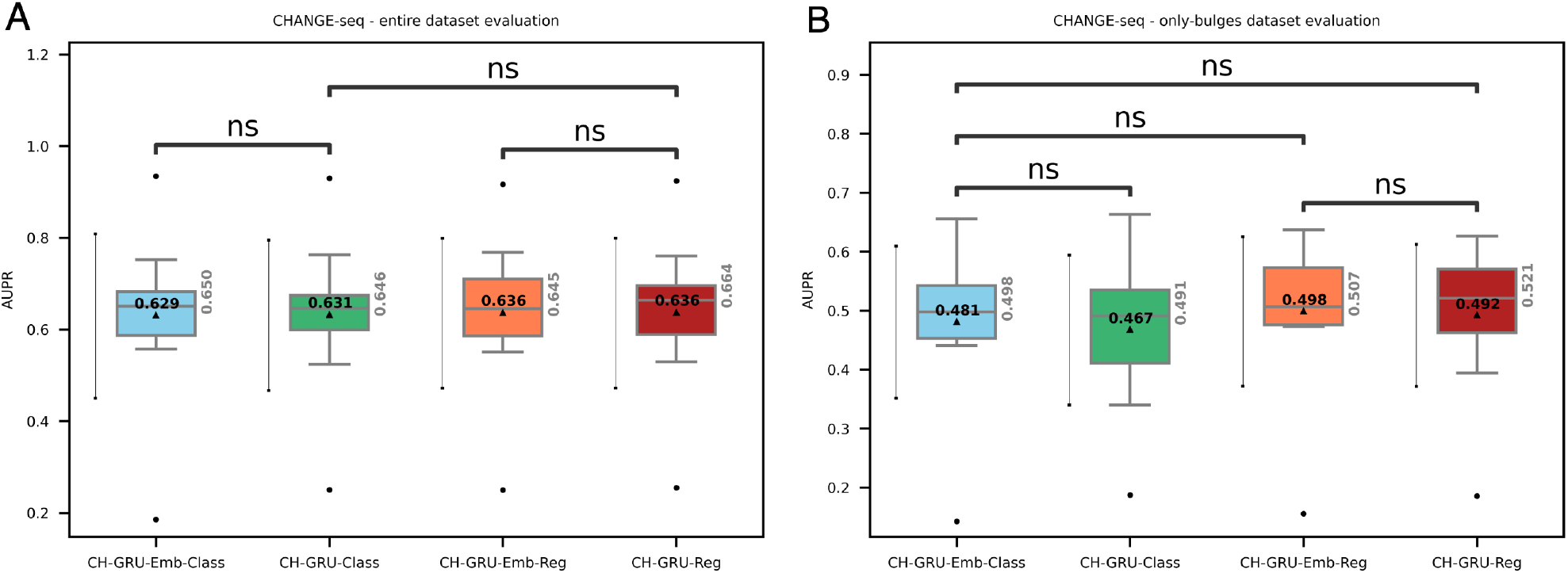
Prediction performance evaluations on *in vitro* data. **(A-B)** A comparison of the CH-GRU and CH-GRU-Emb regression and classification models’ performance on the CHANGE-seq dataset (A) and its only-bulges subset (B). The average AUPR values are denoted by a triangle within each box plot. An error bar of one standard deviation is located on the left of each box plot. The median AUPR values are reported in grey on the right of each box plot. Statistical significance via Wilcoxon signed-rank test is denoted by: ns 5 ·10^*−*2^ < *p* ≤ 1.

**Figure S3.**
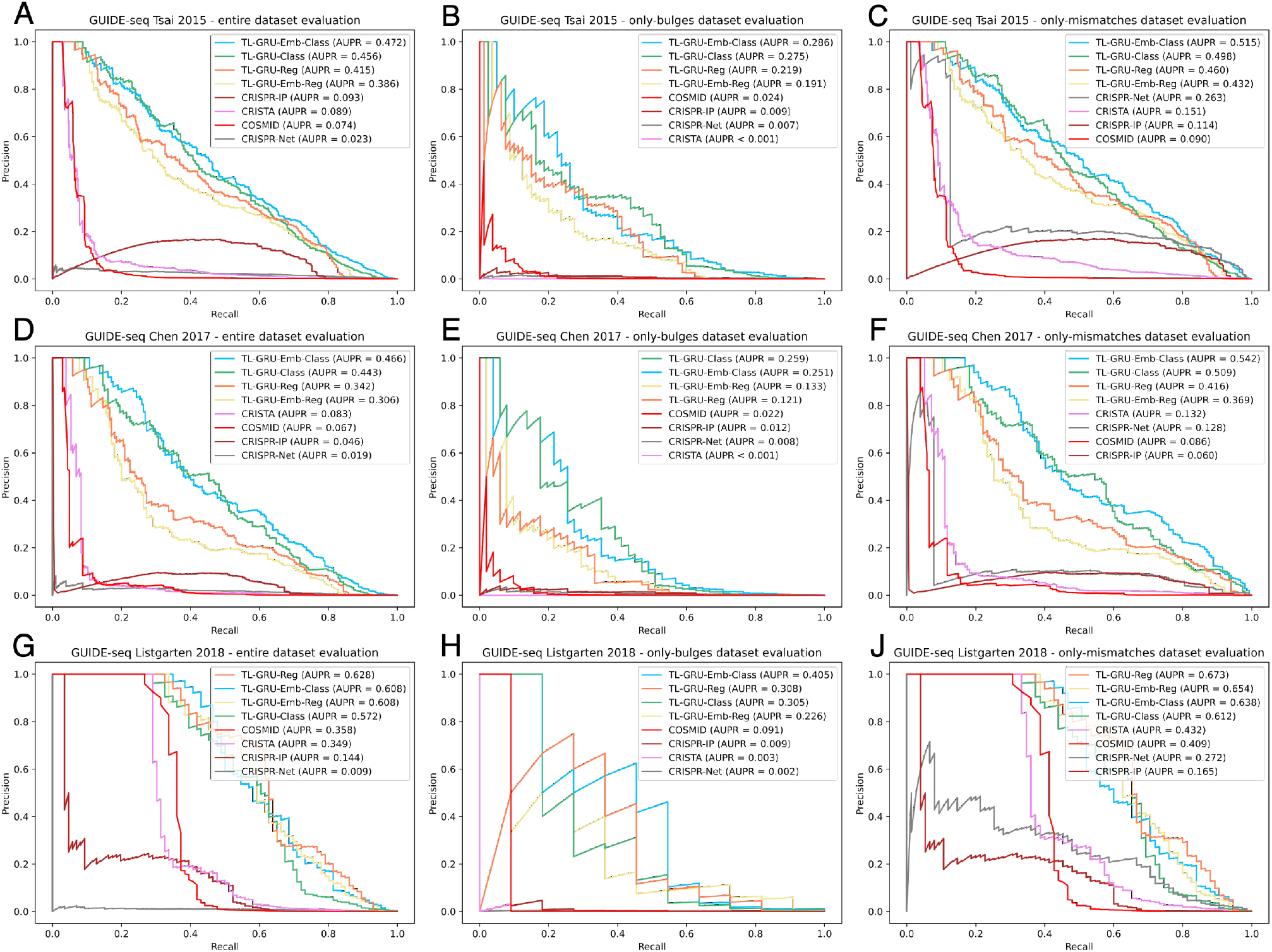
Prediction performance evaluations on independent GUIDE-seq datasets. **(A-J)** Comparison of our models TL-GRU-Reg, TL-GRU-Emb-Reg, TL-GRU-Class, and TL-GRU-Emb-Class with state-of-the-art methods COSMID, CRISTA, CRISPR-Net, and CRISPR-IP on independent GUIDE-seq datasets: Tsai 2015 (A-C), Chen 2017 (D-F), and Listgarten 2018 (G-J). For each dataset, evaluation is performed on the entire dataset (A, D, and G), its only-bulges subset (B, E, and H), and its only-mismatches subset (C, F, and J).

**Figure S4.**
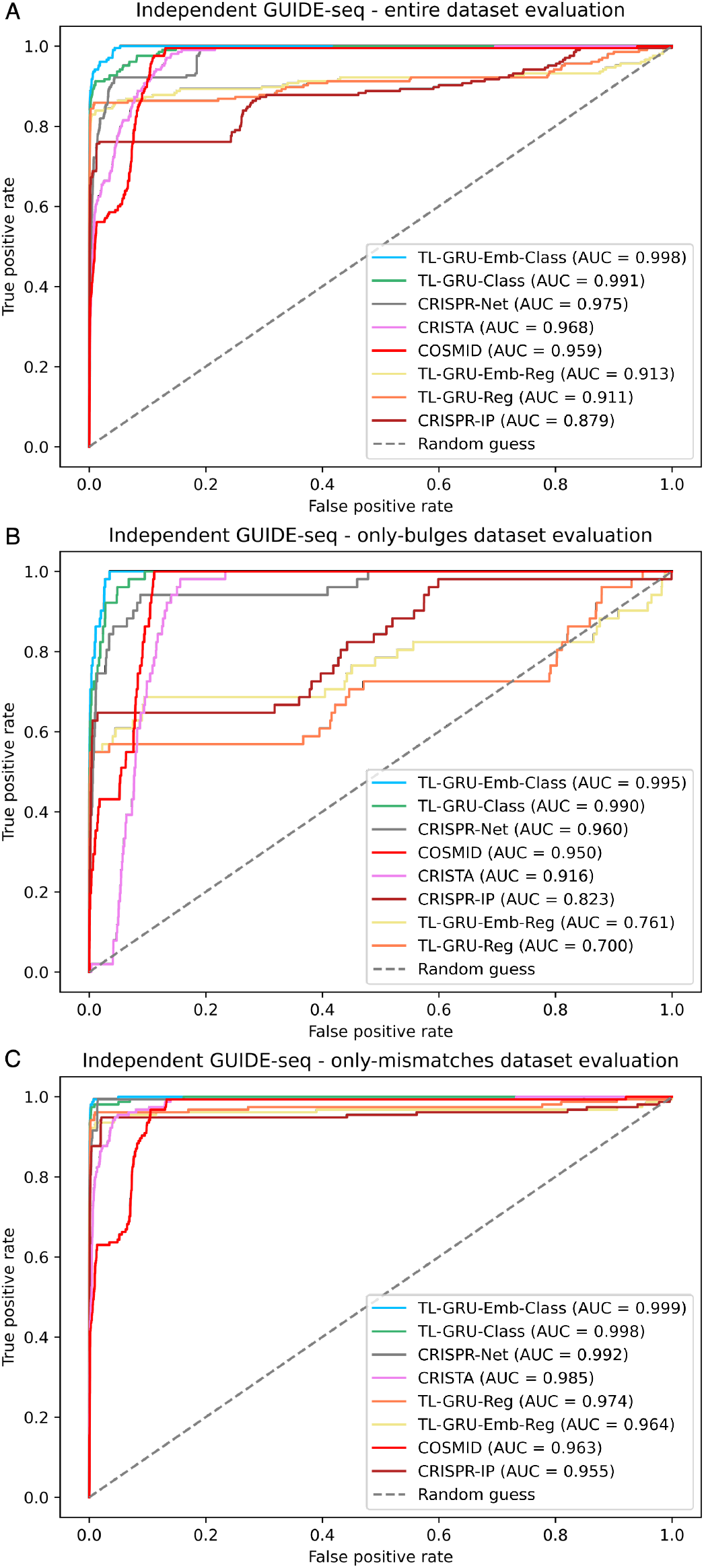
Prediction performance evaluations on the independent GUIDE-seq dataset by the area under the ROC curve. **(A-C)** Comparison of our models TL-GRU-Reg, TL-GRU-Emb-Reg, TL-GRU-Class, and TL-GRU-Emb-Class with state-of-the-art methods COSMID, CRISTA, CRISPR-Net, and CRISPR-IP on our independent GUIDE-seq benchmark dataset (A), its only-bulges subset (B), and its only-mismatches subset (C).

**Figure S5.**
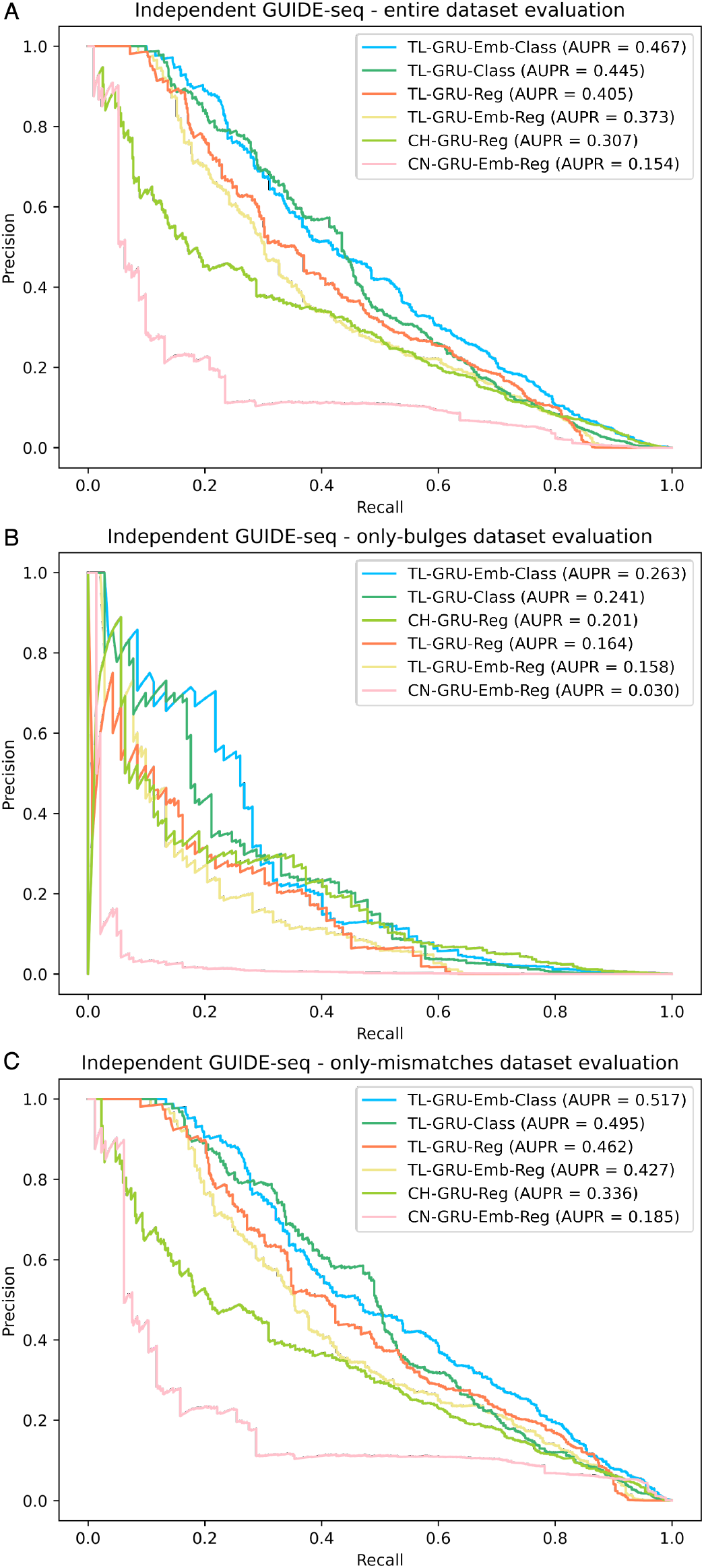
Prediction performance comparison to baseline models on the independent GUIDE-seq dataset. **(A-C)** Comparison of our models TL-GRU-Reg, TL-GRU-Emb-Reg, TL-GRU-Class, and TL-GRU-Emb-Class with top baseline models trained solely on CHANGE-seq (CH) or CRISPR-Net (CN) datasets, CH-GRU-Reg and CN-GRU-Emb-Reg, on our independent GUIDE-seq benchmark dataset (A), its only-bulges subset (B), and its only-mismatches subset (C). The top baseline models for the CH-GRU and CN-GRU models were picked based on the performance on the entire dataset.

**Figure S6.**
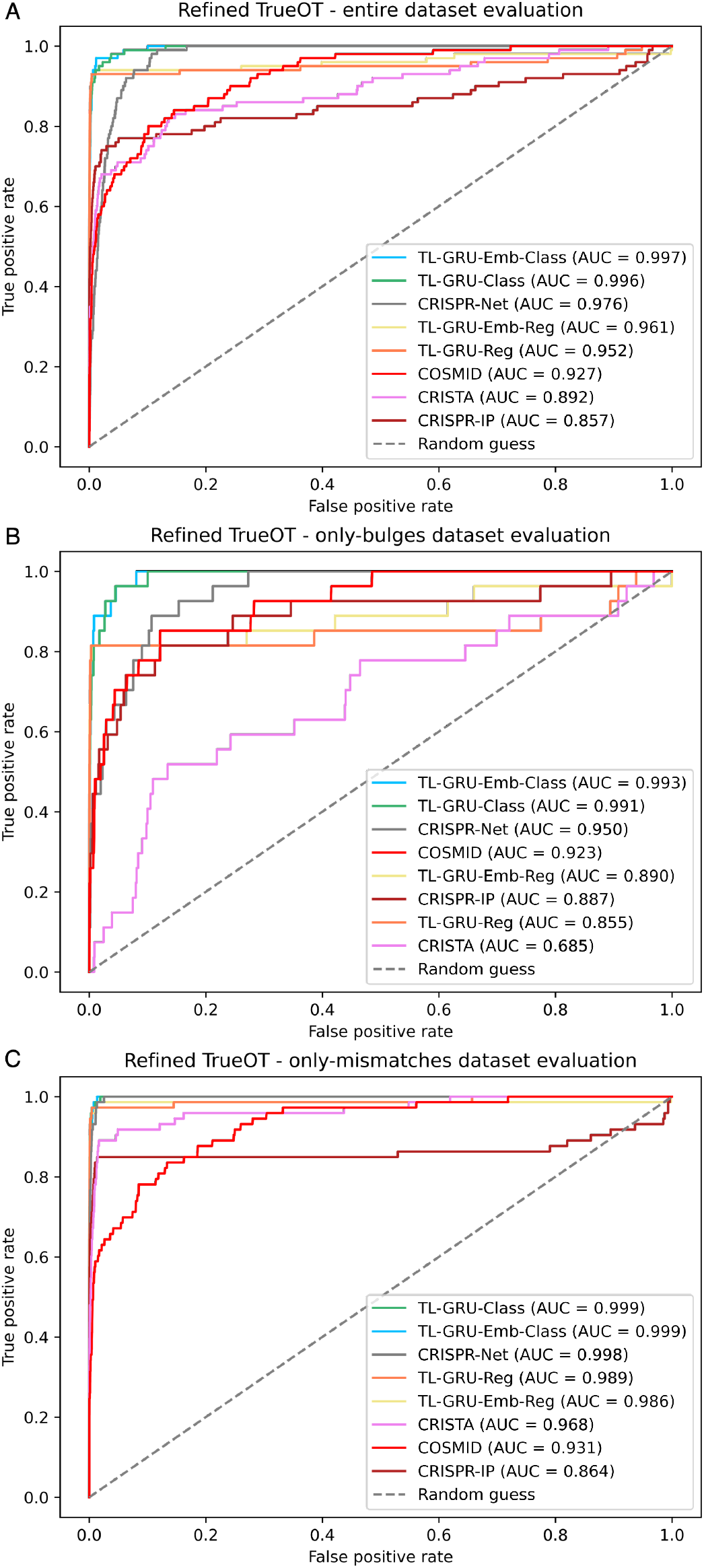
Prediction performance evaluations on the experimentally validated dataset by the area under the ROC curve. **(A-C)** Comparison of our models TL-GRU-Reg, TL-GRU-Emb-Reg, TL-GRU-Class, and TL-GRU-Emb-Class with state-of-the-art methods COSMID, CRISTA, CRISPR-Net, and CRISPR-IP on our refined TrueOT dataset (A), its only-bulges subset (B), and its only-mismatches subset (C).

**Figure S7.**
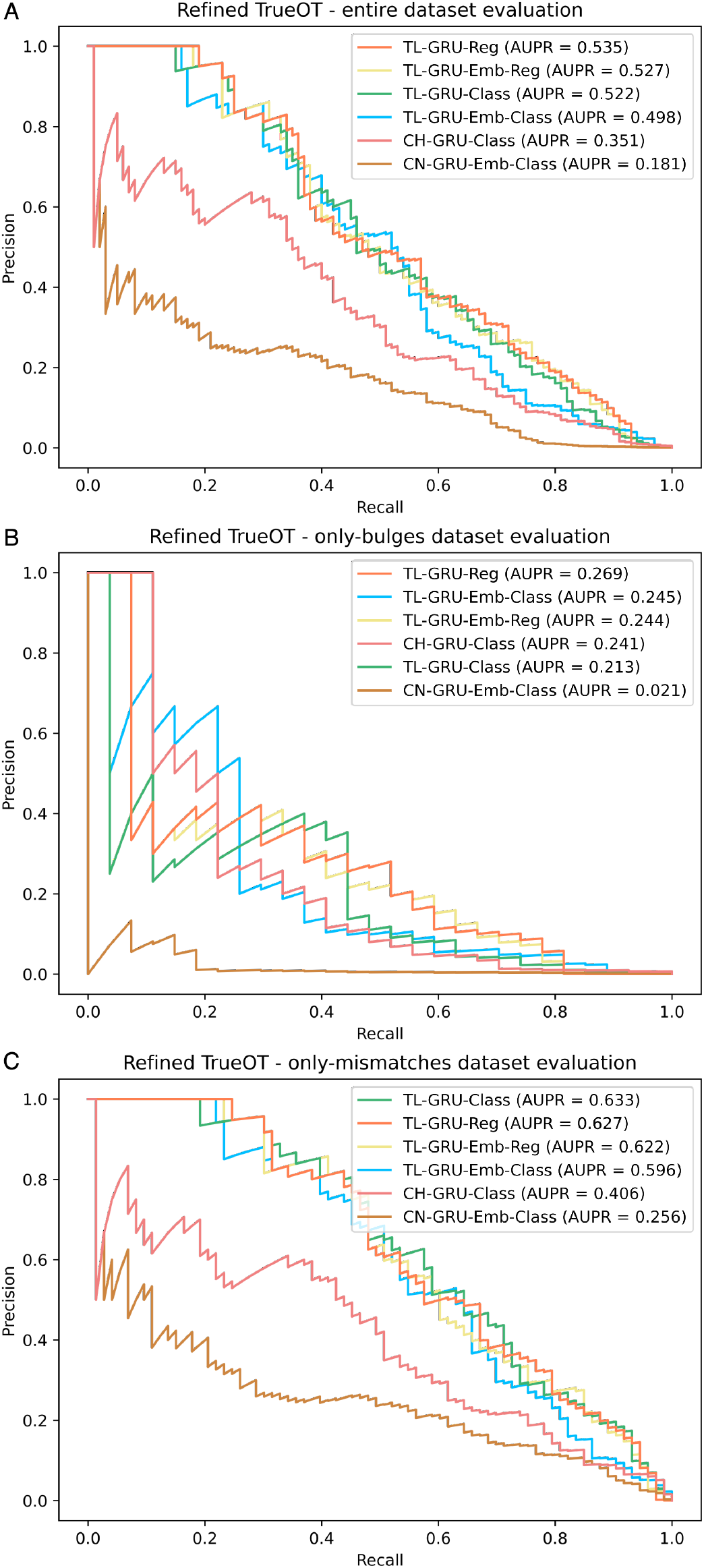
Prediction performance comparison to baseline models on the experimentally validated dataset. **(A-C)** Comparison of our models TL-GRU-Reg, TL-GRU-Emb-Reg, TL-GRU-Class, and TL-GRU-Emb-Class with top baseline models trained solely on CHANGE-seq (CH) or CRISPR-Net (CN) datasets, CH-GRU-Class and CN-GRU-Emb-Class, on our refined TrueOT benchmark dataset (A), its only-bulges subset (B), and its only-mismatches subset (C). The top baseline models for the CH-GRU and CN-GRU models were picked based on the performance on the entire dataset.

**Figure S8.**
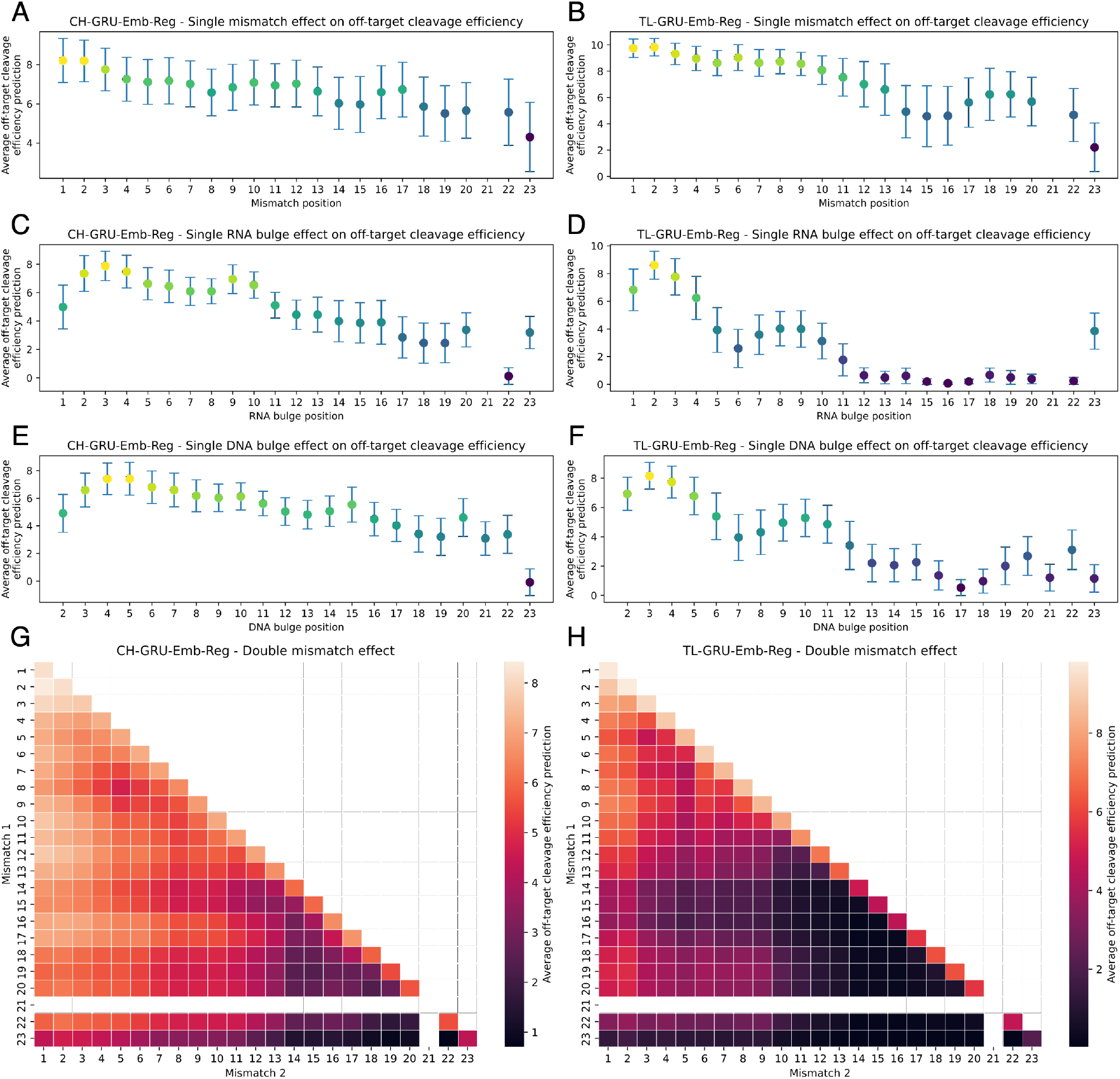
Positional effect of mismatches and bulges on off-target activity learned by our GRU-Emb-Reg model. **A-F** Positional effect of a single mismatch (A-B), RNA bulge (C-D), or DNA bulge (E-F) compared to the sgRNA, predicted by the CH-GRU-Emb-Reg and TL-GRU-Emb-Reg models for *in vitro* (A, C, and E) and *in cellula* (B, D, and F) data, respectively. **G-H** Positional effect of two mismatches compared to the sgRNA predicted by the CH-GRU-Emb-Class (G) and TL-GRU-Emb-Class (H) models.

**Figure S9.**
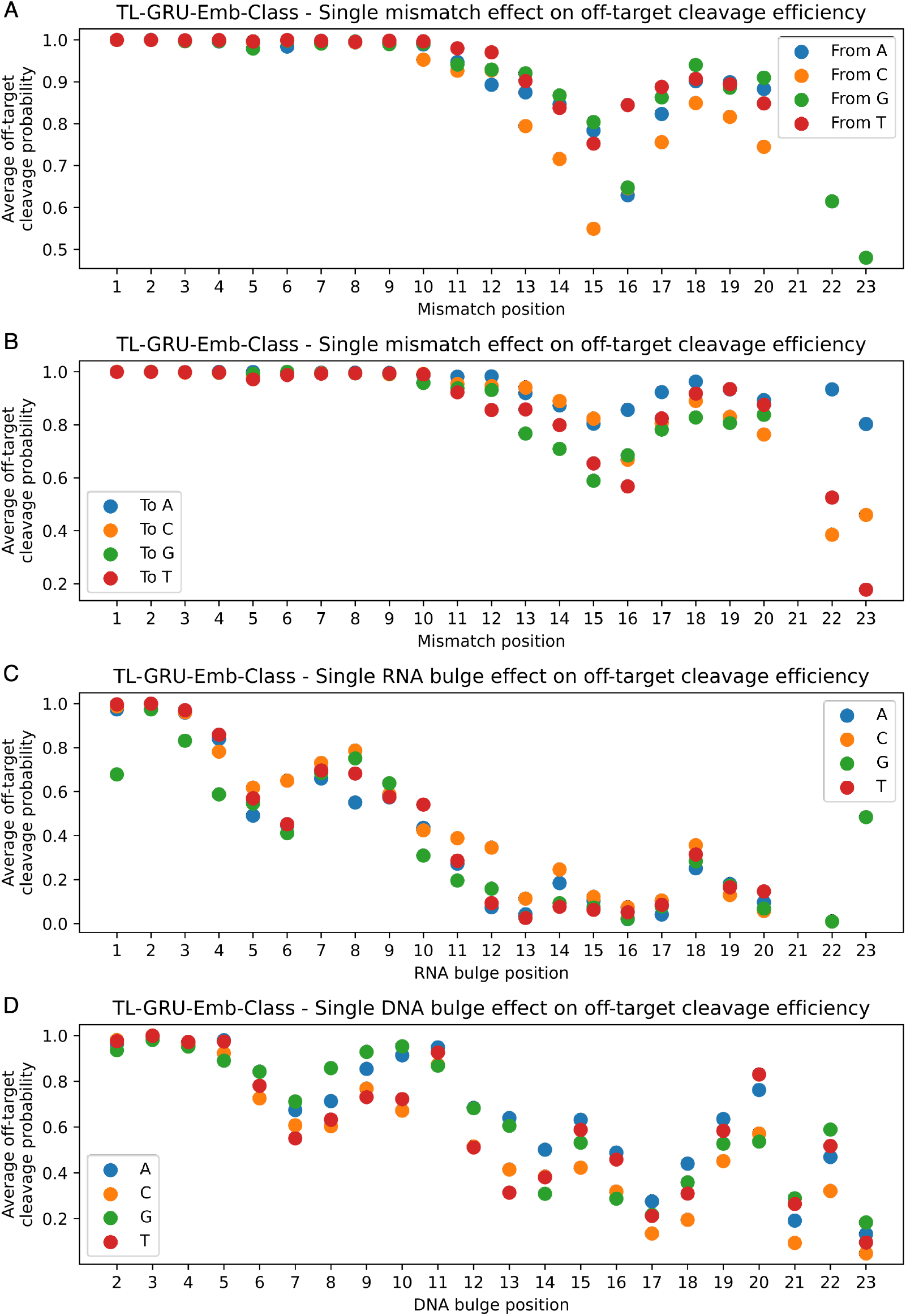
Positional effect of a specific mismatch or DNA/RNA bulge on off-target activity learned by our TL-GRU-Emb-Class model. **A-B** Positional effect of a specific single mismatch at different positions according to the sgRNA (A) and OTS (B) nucleotide. **C-D** Positional effect of a specific RNA (C) or DNA bulge (D).

**Figure S10.**
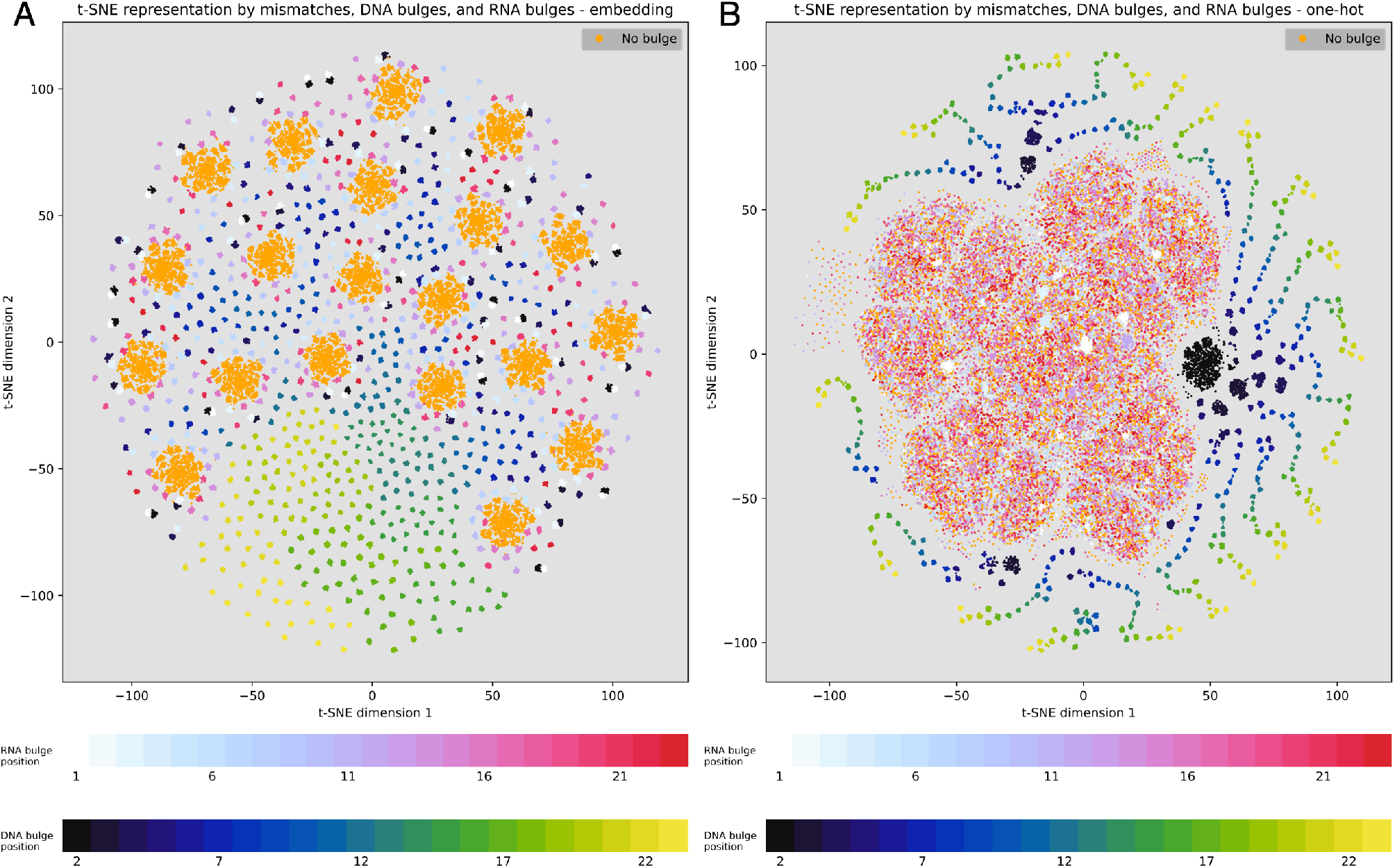
Visualization of the learned embedded representation and one-hot-encoding sequence features of off-target sites with bulges. **A** Learned embedded representation by the TL-GRU-Emb-Class model projected onto a two-dimensional map using t-SNE without subtracting the sgRNA on-target representations. **B** One-hot-encoding sequence features for a pair of aligned OTS and their sgRNAs projected into a two-dimensional map using t-SNE with subtracting the sgRNA on-target one-hot-encoding representations.

